# Opposing roles of the dorsolateral and dorsomedial striatum in the acquisition of skilled action sequencing in rats

**DOI:** 10.1101/2021.04.15.439935

**Authors:** Karly M. Turner, Anna Svegborn, Mia Langguth, Colin McKenzie, Trevor W. Robbins

## Abstract

The shift in control from dorsomedial to dorsolateral striatum during skill and habit formation has been well established, but whether striatal subregions orchestrate this shift co-operatively or competitively remains unclear. Cortical inputs have also been implicated in the shift towards automaticity, but it is unknown if they mirror their downstream striatal targets across this transition. We addressed these questions using a five-step heterogeneous action sequencing task in rats that is optimally performed by automated chains of actions. By optimising automatic habitual responding, we discovered that loss of function in the dorsomedial striatum accelerated sequence acquisition. In contrast, loss of function in the dorsolateral striatum impeded acquisition of sequencing, demonstrating functional opposition within the striatum. Unexpectedly the medial prefrontal cortex was not involved, however the lateral orbitofrontal cortex was critical. These results shift current theories about striatal control of behavior to a model of competitive opposition, where the dorsomedial striatum acts in a gating role to inhibit dorsolateral-striatum driven behavior.

## INTRODUCTION

There is mixed consensus on exactly how habits and skills interact. They are two separate descriptors of behavior with overlapping and distinct features (Dezfouli & Balleine, 2012; Graybiel & Grafton, 2015; Jin & Costa, 2015; Robbins & Costa, 2017). Skills typically describe refined behavioral repertoires, which may be under goal-directed or habitual control (or a combination as described by hierarchical accounts). In contrast, habits are defined as responses that are triggered by stimuli and are autonomous of the outcome value but may include both skilled and unskilled behaviors. The concept of automaticity captures many of the shared elements between habits and skills, where the behavior becomes stereotypical, performed with little variation in a highly efficient manner and without effortful thought (Ashby, Turner, & Horvitz, 2010). Chunked action sequences provide an opportunity to study the nexus of automaticity, skills, and habits (Dezfouli, Lingawi, & Balleine, 2014; Graybiel & Grafton, 2015; Robbins & Costa, 2017). The transition to automaticity in both habits and skills is paralleled by a well-documented shift in control from the dorsomedial (DMS) to dorsolateral (DLS) striatum (Ashby et al., 2010; Graybiel & Grafton, 2015; Kupferschmidt, Juczewski, Cui, Johnson, & Lovinger, 2017; Thorn, Atallah, Howe, & Graybiel, 2010; Yin et al., 2009). Yet, it is unknown how this transition occurs and how these regions co-ordinate the control of actions (Bergstrom et al., 2018; Kupferschmidt et al., 2017).

Goal-directed behavior, dependent on DMS function, dominates early in instrumental conditioning but if conditions support habitual responding then the DLS takes control (Balleine, Liljeholm, & Ostlund, 2009; Yin & Knowlton, 2006; Yin, Knowlton, & Balleine, 2004, 2005, 2006; Yin, Ostlund, Knowlton, & Balleine, 2005). Similarly, in skill learning there is an early learning phase where actions are variable and slow but as they become refined and efficient then control shifts from the DMS to DLS (Kupferschmidt et al., 2017; Lehericy et al., 2005; Miyachi, Hikosaka, & Lu, 2002; Yin et al., 2009). Neural studies indicate the DMS and DLS operate in parallel during this transition with some degree of interdependency (Gremel & Costa, 2013; Vandaele et al., 2019; Yin et al., 2009). More recently it has been shown that the DLS is engaged from the beginning of conditioning and only after initial experience does the goal-directed system start driving behavior (Bergstrom et al., 2018; Kupferschmidt et al., 2017; A. C. W. Smith et al., 2021). However, it is unclear whether the DMS and DLS act via a co-operative or competitive relationship (Balleine et al., 2009; K. S. Smith & Graybiel, 2016). Dual control accounts suggest these two processes both contribute to behavior with the relative influence shifting with extended training (Balleine, 2019; Balleine & Dezfouli, 2019; Dickinson, 1985; Perez & Dickinson, 2020; Robbins & Costa, 2017). It was recently proposed that responses reflect the summation of goal-directed and habitual processes (Perez & Dickinson, 2020). Alternatively, habits may form early but remain latent or inhibited unless required (Hardwick, Forrence, Krakauer, & Haith, 2019). Similar accounts may apply to the relative neural contribution of the DMS and DLS to action control.

If these regions operate independently, then loss of function should impair only that region’s function (e.g., loss of DMS leads to impaired goal-directed action), however if they operate co-operatively or co-dependently then suboptimal performance would be expected in both functions (e.g. loss of DMS also impairs habit formation). In contrast, an opponent relationship would predict that loss of function in one region would favour the alternate structure’s function (e.g., loss of DMS leads to *enhanced* habit formation). The role of cortical inputs may be critical in modulating this striatal balance (Daw, Niv, & Dayan, 2005; Peak, Hart, & Balleine, 2019). A problematic issue when addressing this question in habits has been the "zero-sum" interpretation as habits are defined by a lack of goal-directed features (Balleine & Dezfouli, 2019; Robbins & Costa, 2017; Schreiner, Renteria, & Gremel, 2020). However, a loss of devaluation sensitivity may result from impaired instrumental learning, rather than habit formation (Balleine & Dezfouli, 2019). Habits are typically identified by an impairment in action modification when conditions change (e.g., devaluation or contingency degradation), and rarely as the optimal response in a task. To address this issue, it was recently suggested habits can be defined by four features: rapid execution, invariant response topography, action chunking, and insensitivity to outcome value and contingency (Balleine & Dezfouli, 2019). Hence, we developed a novel rodent paradigm using a sequence of heterogeneous actions where automated, reflexive responding would lead to superior performance, to test models of striatal control during the development of automaticity. We hypothesised that DMS loss of function would causally accelerate, whereas DLS loss of function would impair, the development of behavioral automaticity.

## RESULTS

### A novel five-step action sequencing task for rats

Using a multiple-response operant chamber (Carli, Robbins, Evenden, & Everitt, 1983), rats made a nose poke response in each of five holes from left to right to receive a reward sucrose pellet in the magazine (Figure 1H). After brief training, rats could initiate self-paced sequences during a daily 30 min session (Figure 1G). Importantly, the sequential nose poke task was self-initiated and not cued. This required the acquisition and then retrieval of a planned motor sequence, of which the first four actions were never immediately rewarded. The removal of cues also ensured that the sequence required internal representation where enhanced performance was due to an improved representation and retrieval of the sequence rather than an improved ability to detect stimuli (Yin, 2010). It was expected that following repeated reinforcement the five individual actions would be chunked into a more efficient unitary motor program. This task would be most efficiently performed by the development of automaticity and aimed to fit the behavioral criteria for both habits and skills.

**Figure 1.**
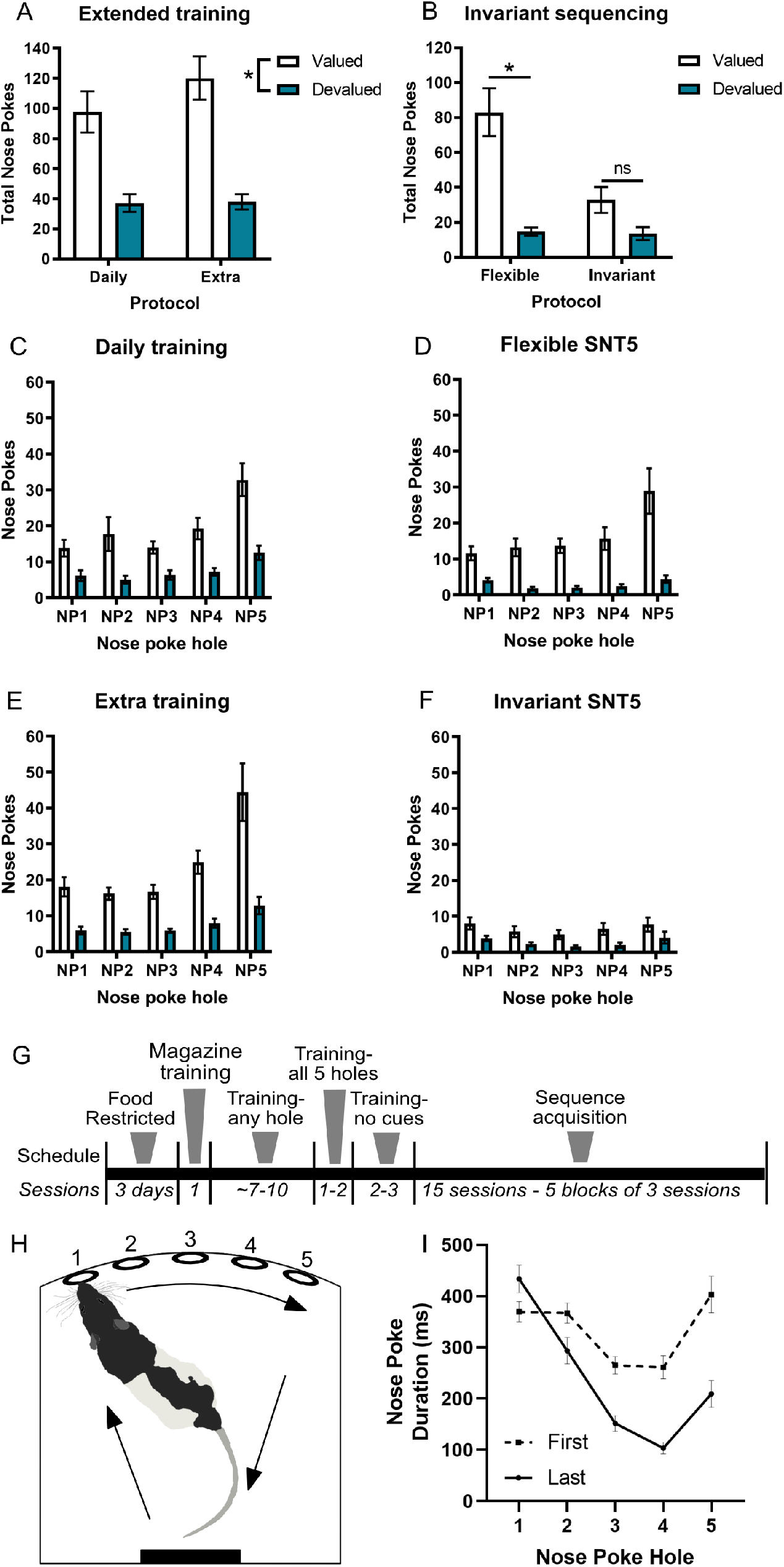
The sequential nose poke task leads to ballistic responding. (A) Extended training did not alter sensitivity to outcome-specific devaluation between the daily and extended training groups, with both groups responding more in the valued compared to devalued test session. (C) and (E) show the number of responses across the sequence elements. (B) Constraining rewards to only perfect sequences with time-outs for any errors in the invariant protocol led to habitual responding, while the flexible group remained goal directed. (D) and (F) show the number of responses across the sequence elements under the valued and devalued conditions. (G) The training schedule included habituation to the magazine and nose poke training. The nose poke cues were rapidly removed once rats were responding to each hole. From the beginning of the sequence acquisition period only correct five-step sequences were rewarded and errors were penalised by a brief time out period, after which the sequence had to be reinitiated (see Methods). (H) Rats were trained to make a five-step nose poke sequences to receive a food reward if they nose poked into each of the holes in order from left to right. (I) Rats developed a ballistic response pattern across the five holes from the first to last block of training. Each nose poke was faster and with less variance as the sequence progressed. Data shown as group mean ± S.E.M. *p<0.05.

### Testing for habitual properties of the heterogenous 5-element response sequence

A classic method used to induce habitual responding is extended training (Dickinson, 1985). We trained rats to perform the sequencing task without cues and then placed half of them onto a twice daily (extended) training regime, while the other half continued with daily sessions for 10 days. Outcome-specific devaluation was then used to probe habits through sensitivity to changes in outcome value. The outcome was devalued by providing free access to 25 g of the sucrose pellets, allowing the rat to become sated, before recording sequencing responses for 10 min in extinction. This was compared to a separate counterbalanced session where rats were sated on grain pellets before testing, thereby leaving the outcome (sucrose pellets) still valued. Rats were tested in extinction to prevent learning about the change in outcome value through the experience of earning the outcome in the sated state, thereby demonstrating whether actions were influenced by changes in inferred outcome value. If the rats respond less when sated on sucrose pellets than grain pellets, then the specific value of the outcome was being used to adapt actions and the animal was responding under goal-directed control. If the rat responded equally after both the sucrose and grain pellets, then changes in outcome value were not being used to guide actions, indicative of habits. There was no evidence of habit formation in either group with a significant effect of Devaluation (F_1,22_=67.78, p<0.001) and Hole (F_2,38_=29.66, p<0.001), but no main effect of Group (F_1,22_=0.9, p=0.4) or interactions with Group (p’s>0.3) (Figures 1A, C, E). This indicates both groups remained goal-directed.

Another important factor in habit formation is behavioral variation (Dickinson, 1985). If rats made a sequencing mistake (most commonly skipping a hole due to insufficient nose poke depth) the program would wait for the correct response before moving to the next hole. This allowed rats to correct their mistakes and then continue with the sequence. This resulted in occasional variation in rewarded sequence structure (e.g., **<u>1</u>**-**<u>2</u>**-4-2-**<u>3</u>**-**<u>4</u>**-**<u>5</u>**-**<u>reward</u>**) and promotes some level of self-monitoring to detect where an error was made so it could quickly be rectified. Variation in sequence structure and attending to actions to detect errors should retard habit formation. Given there were no differences detected after extended training, the rats were again split into two groups (n=12/group, balanced for prior training) with one group moving to an invariant sequencing protocol where errors were punished, and the other group continued training on the same protocol. In the new, invariant protocol, when rats made a sequencing error the house light was illuminated for 5 s and they then needed to restart the sequence from the beginning, ensuring only perfect sequences were rewarded (e.g., **<u>1</u>**-**<u>2</u>**-4- time out-**<u>1</u>**-**<u>2</u>**-**<u>3</u>**-**<u>4</u>**-**<u>5</u>**-**<u>reward</u>**). Rats were then retested for habitual responding using outcome-specific devaluation as described above. Across the five holes, there was a main effect of Devaluation (F_1,22_=38.57, p<0.001) and Hole (F_1,30_=9.39, p=0.002) and Group (F_1,22_=8.19, p=0.009) and Devaluation X Hole X Group interaction (F_4,88_=6.95, p<0.001) (Figures 1B, D, F). Importantly there was a significant Devaluation X Group interaction (F_1,22_=12.11, p=0.002), demonstrating devaluation sensitivity was significantly reduced when only perfect sequences were rewarded. A simple effects test revealed that while the flexible group remained goal-directed (p<0.001), the invariant protocol led to habitual responding as indicated by the lack of a significant difference in responding between the valued and devalued conditions (p=0.07). There was a clear reduction in the amount of valued responding with the introduction of the invariant procedure compared to the flexible group (Figure 1B). We considered if this reduction in responding during the valued session could be due to poor goal-directed learning, but the chance of producing 5 uncued actions in the correct order without knowledge of the action-outcome association during training is highly unlikely. This is thus the first demonstration of habitual responding on a heterogenous action sequencing task in rodents and all subsequent experiments in this study used this version of the task.

Unfortunately rats rapidly ceased responding under extinction conditions, producing very few complete sequences, which was not unexpected given the sequencing task uses a continuous reinforcement schedule (see Figure 1F noting five nose pokes were required per sequence). This led to floor effects for measuring sequencing behavior (such as timing or effects on initiation, execution, and terminal elements) and was likely to restore goal-directed control very quickly when extinction was detected. The significant 3-way interaction indicated that the flexible group showed outcome devaluation sensitivity on every hole (p<0.001), whereas the invariant group responded habitually on nose pokes 2, 4 and 5 (p’s>0.08) but showed outcome sensitivity on nose pokes 1 (p=0.02) and 3 (p=0.04).

However, there was no significant difference in the frequency of nose pokes across holes 1-5 within either the valued or devalued session for the invariant group, indicating that rats did not perform the initiation, execution, or termination elements significantly more under either condition. They reduced responding across all holes, indicative of the sequence becoming chunked into a single motor plan that was no longer under goal-directed control. Although devaluation is the ’gold-standard’ test for detecting habits, the lack of whole sequences performed and limited scope for reliably detecting differences between experimental groups where smaller effect sizes were expected, prevented its use in subsequent experiments.

In a separate cohort of rats, we then measured hallmark traits of automaticity - increased speed and reduced variability. Acquisition of sequencing was observed over 15 sessions that were grouped into five blocks of three sessions (Figure 1G). Response times across the five actions in the first acquisition block were comparable but following further training, a ballistic response pattern developed. This pattern was characterised by an extended initiation pause prior to the first element that led into a rapid escalating response pattern from holes 2-4, being completed with a concatenation pause following the terminal element. Here the rat anticipates and prepares the next motor chunk - reward retrieval. Data from treatment-naïve rats (n=36) trained on the finalised version of the sequencing task (see Figure 1G), indicated that from the first to last block there was a significant change in nose poke duration at each location (interaction F_2,82_=19.80, p<0.001; pairwise comparisons p’s<0.025) (Figure 1I). The ballistic response pattern began to emerge in the first block with relatively equivalent variation at each step. By the last block, each action in the sequence became increasingly faster and less variable, indicative of refined and automated action sequencing. This response pattern, particularly the initiation and termination delays, are characteristic of motor sequence chunking (Abrahamse, Ruitenberg, de Kleine, & Verwey, 2013; Sternberg, Monsell, Knoll, & Wright, 1978). Therefore, the sequential nose poke task leads to chunked action sequencing with features of both skill and habit formation as defined by rapid execution, invariant response pattern, evidence of sequence chunking and insensitivity to changes in outcome value (Balleine & Dezfouli, 2019).

### DMS-lesioning improved acquisition of action sequencing, while DLS-lesioning impaired efficient sequencing

#### Initial training

To determine if the DMS and DLS work cooperatively or in opposition, subregion- specific loss of function was required throughout training and 15 sessions of sequence acquisition (Figure 2A, B). Lesions made via discrete fiber-sparing quinolinic acid infusions avoided any overlap between the DMS and DLS. Following recovery, rats were food restricted and trained on the sequencing task (see Figure 1C for schedule). Our a priori hypothesis was that we would observe divergence between DMS and DLS groups and hence direct comparisons were made. DLS-lesioned rats took significantly longer to reach training criteria (Figure 2C; Lesion: F_2,23_=7.80, p=0.003) than DMS-lesioned (p=0.045) or sham treated rats (p=0.001). Rats then moved to sequence acquisition where only perfect 5-step sequences were rewarded.

**Figure 2.**
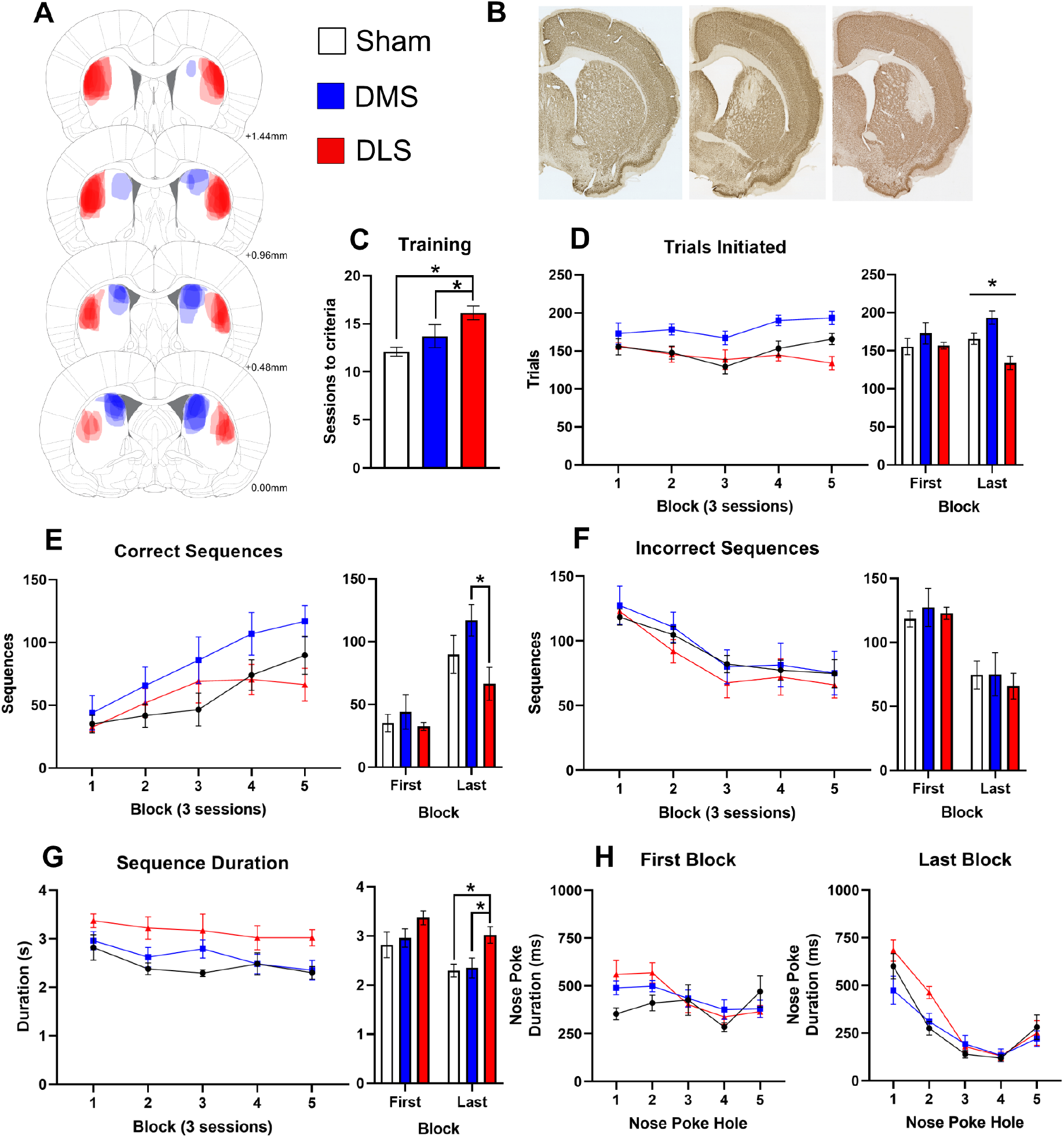
DMS-lesioning improved acquisition of action sequencing, while DLS lesioning impaired efficient sequencing. (A) Rats received targeted bilateral lesions with extent illustrated for lesion groups; sham (open, n=11), DMS (blue, n=7), DLS (red, n=8). (B) Striatal sections showing NeuN staining in sham (left), DMS (middle) and DLS (right) lesioned rats. (C) DLS-lesioned rats required significantly more sessions to reach training criteria than sham or DMS-lesioned rats. (D) Left: When acquiring sequencing behavior, DMS-lesioned rats initiated more trials than either DLS-lesioned or sham rats. Right: There was no significant difference between groups in the first block, however by the last block, DLS-lesioned rats started fewer trials and DMS-lesioned rats completed more trials than sham. (E) Left: Contrasting effects of lesions were also observed for the number of correct sequences. Right: DMS-lesioned rats completed nearly twice as many correct sequences than DLS-lesioned rats in the last block of acquisition. (F) Incorrect sequences decreased across acquisition, demonstrating all groups learned to avoid errors. (G) Left: Overall, DLS-lesioned rats took longer to complete sequences than sham rats. Right: All groups completed sequences significantly faster from the first to last block of acquisition and in the final block DLS-lesioned rats took significantly longer to complete sequences than sham and DMS-lesioned rats. (H) Across acquisition, the duration of nose pokes became faster and developed a ballistic response pattern. Right: By the last block, DLS-lesioned rats paused significantly longer than DMS-lesioned rats on the first two actions of the sequence, but not the latter half of the sequence. Data shown as group mean ± S.E.M. *p<0.05.

#### Sequence acquisition

We compared performance measures during acquisition to quantify action sequence refinement, with a focus on changes between the first (sessions 1-3) and last blocks (sessions 12-15). Across acquisition, DMS-lesioned rats initiated more trials (Figure 2D; Lesion: F_2,23_=6.94, p=0.004) than either DLS-lesioned (p=0.002) or sham control (p=0.005) rats. The number of trials initiated was equivalent between groups in the first block (p>0.3). However, by the last block there were opposing effects detected between groups (Lesion: F_2,23_=11.59, p<0.001). DLS-lesioned rats initiated significantly fewer trials than sham rats (p=0.09), while DMS-lesioned rats completed significantly more trials than sham rats (p=0.025); with a substantial difference between DMS and DLS groups (p<0.001). This acquired divergence between DMS and DLS lesioned rats demonstrated that DMS-lesions enhanced, while DLS- lesions impaired, initiation of action sequences. As sessions were time limited, performing more trials indicated greater speed and opportunity for reward, however these trials could have been either correct or incorrect.

The number of correct sequences increased for all groups across acquisition, indicating all groups were able to learn the five-step sequence. Opposing effects of striatal lesions were again also observed in the total number of correct sequences. There was no difference between groups on the first block, yet there was a clear divergence between DMS and DLS lesioned rats across acquisition (Figure 2E). Our a priori hypothesis was that DMS and DLS lesions would have opposing effects and a comparison between these lesion groups found that DMS-lesioned rats completed nearly twice as many correct sequences as DLS-lesioned rats at the end of acquisition (DMS = 117±12, DLS = 67±13; t_13_=2.79, p=0.015).

Despite the dissociation between groups in both the number of sequences initiated and correct sequences, there was no difference in the number of incorrect sequences made by each group (Figure 2F; F_2,23_=0.16, p=0.85) and all groups showed a significant reduction in erroneous sequences from the first to last block (p’s<0.01). These results support a model of sequence learning where the DMS and DLS have opposing roles in the development of automated behaviors.

#### Sequence timing

We next investigated how striatal lesions influenced the timing of actions within sequences. Across sequence acquisition, sequence duration significantly reduced (Figure 2G; F_4,92_=6.74, p<0.001), indicating increased sequencing efficiency with experience. This is important as faster execution is considered one of the hallmarks of skill learning and sequence chunking. Throughout acquisition, DLS-lesioned rats took significantly longer to execute complete sequences (Lesion: F_2,23_=4.59, p=0.021) than sham rats (p=0.007), with a trend towards impairment compared to DMS-lesioned rats (p=0.059). All groups completed sequences significantly faster from the first to last block of acquisition (sham, t_10_=2.33, p=0.042; DMS, t_6_=4.78, p=0.003; DLS, t_7_=2.83, p=0.026). In the final block, DLS-lesioned rats took significantly longer to complete sequences (Lesion: F_2,23_=5.87, p=0.009) than sham (p=0.004) and DMS-lesioned rats (p=0.013), supporting the conclusion that DLS lesions impaired the development of refined action sequencing. In addition, we examined each individual rat’s standard deviation of sequence duration to determine if variability reduced with training, as another hallmark of skill learning and automaticity. Only the DMS-lesioned rats had a significant reduction from the first to last block in their individual sequence duration variability (sham p=0.95; DMS p=0.021; DLS p=0.16), in agreement with other measures indicating enhanced automatisation of sequencing with DMS lesions (Supplementary Figure 1C).

As the task utilised five spatially heterogeneous responses, the timing of each action within the sequence was then compared across the initiation (hole 1), execution (holes 2-4) and terminal (hole 5) responses as well as the nose poke duration within each hole. Nose poke duration became faster across acquisition (Figure 2H; Block: F_4, 92_=19.57, p<0.001) and developed the characteristic accelerating response pattern (Block X Hole: F_16, 368_=9.07, p<0.001). There were no significant differences between groups in the first block of acquisition (Figure 2H; F_2,23_=0.67, p=0.52). However, by the last block, nose poke duration had stabilised to a ballistic response pattern and the variance in timing had reduced as the movement became stereotypical. On the last block, there was a main effect of Hole (F_2,49_=67.84, p<0.001) and the Hole X Lesion interaction approached statistical significance (F_4,49_=2.51, p=0.051). Planned post-hoc comparisons found DLS-lesioned rats paused significantly longer than DMS-lesioned rats on the first two actions of the sequence (hole 1, t_13_=2.28, p=0.040 and hole 2, t_13_=2.92, p=0.012) but not the latter half of the sequence (p’s>0.7). These results demonstrated that while DLS-lesioned rats were capable of extremely fast nose poke responses (see hole 4) and therefore were not exhibiting general motor impairments (also see locomotion data in Supplementary Figure 1G), they were significantly delayed in initiating the sequence. These results indicated that the DLS is important for action selection or retrieval. However, once the sequence was engaged, its execution was not dependent on intact DLS function.

The inter-poke interval between correct nose pokes also speeded with training (Supplementary Figure 1A, B) indicating improved efficiency. There was a u-shaped pattern across the curved wall, likely reflecting ambulation requirements (F_3, 45_=44.88, p<0.001) but there was no effect of lesion (F_2, 23_=1.04, p=0.37) or interactions with treatment groups (p’s>0.1). There was also no effect of group on the latency from leaving the magazine to starting at hole 1 (Supplementary Figure 1D), indicating all groups were equally as motivated to initiate sequences. There were also no significant changes in magazine nose poke duration (Supplementary Figure 1F) or reward collection latency (Supplementary Figure 1E) over acquisition or between groups suggesting training and lesions did not alter reward motivation.

Across numerous measures of performance, our results showed that DMS lesions accelerate the shift towards automatisation, while DLS lesions impair the development of efficient action sequencing. Delayed sequence initiation but not execution or termination in DLS-lesioned rats, suggest that the DLS is important for loading the motor program, but once rats started responding the transition between elements was accurate and rapid, indicative of action sequence chunking. Cortical inputs to the striatum play an important role in both adaptive and habitual responding therefore we sought to determine whether subregions within the prefrontal cortex influence the acquisition of action sequencing. We hypothesised that cortical regions with inputs into the DLS would impair sequence acquisition, while those with inputs to the DMS may enhance acquisition.

### Lateral OFC but not medial OFC lesions impair sequencing

We first examined the role of the medial (mOFC) and lateral (lOFC) orbitofrontal cortex, which project to medial and lateral regions of the dorsal striatum, respectively. mOFC lesions lead to habitual responding via an inability to retrieve outcome value in outcome devaluation tests (Bradfield, Dezfouli, van Holstein, Chieng, & Balleine, 2015; Bradfield, Hart, & Balleine, 2018). In contrast, the lOFC is well known for its role in flexible responding in reversal learning, outcome prediction and devaluation (Gremel et al., 2016; Gremel & Costa, 2013; Hervig et al., 2019; Izquierdo, 2017; Panayi & Killcross, 2014; Turner & Parkes, 2020). However, it was unclear whether these regions would enhance action sequencing.

#### Acquisition of sequencing

Using the same procedure, we determined if the mOFC and lOFC were required for action sequencing (Figure 3A, B). There was no effect of lesions on the number of sessions required during training (Figure 3C). Across sequence acquisition, there was a significant interaction between groups with lOFC-lesioned rats starting significantly fewer trials than the mOFC group in the final two blocks (Figure 3D; Lesion X Block, F_8, 180_=2.72, p=0.024).

**Figure 3.**
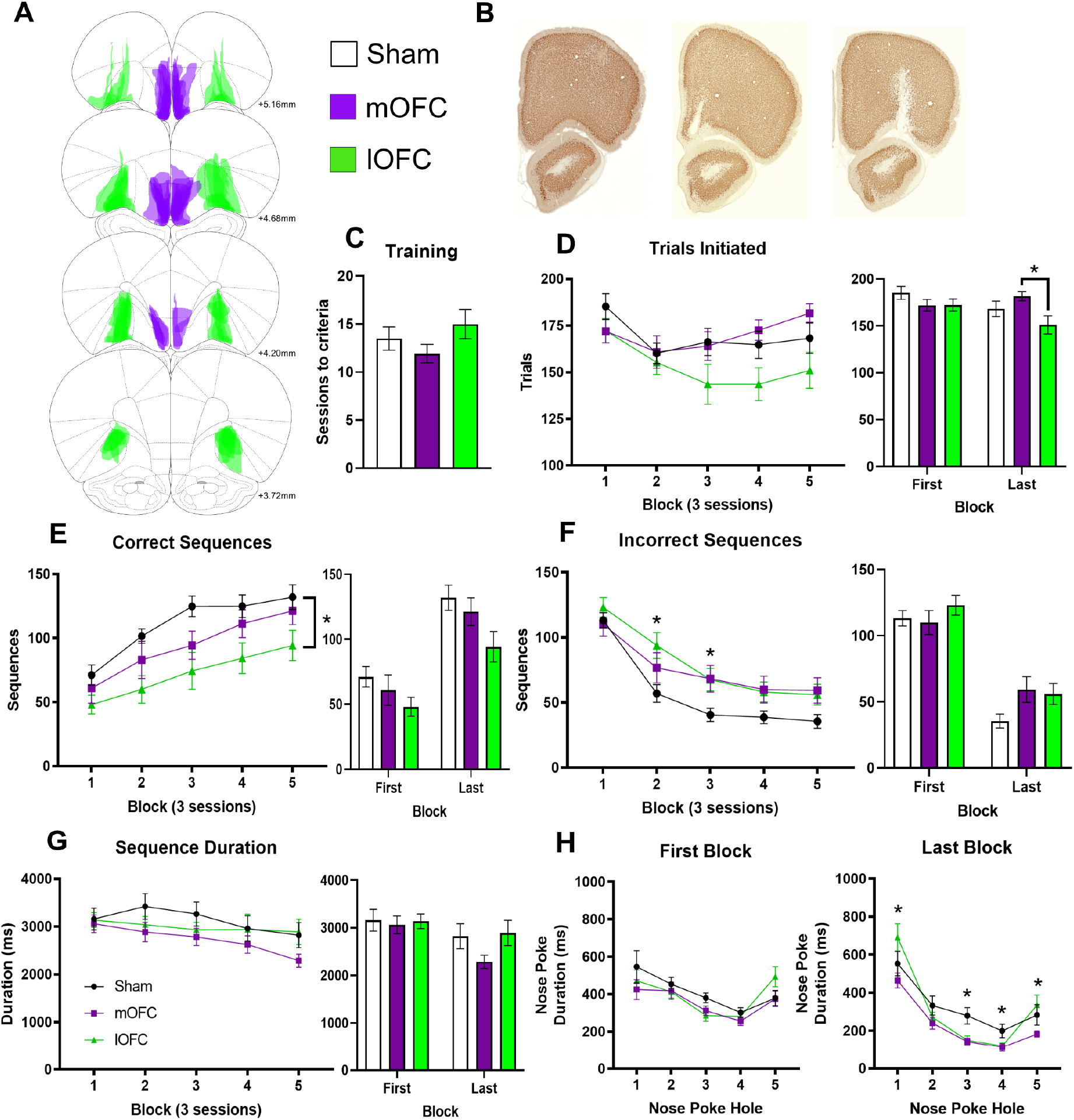
Lateral OFC but not medial OFC lesions impair sequencing. (A) Rats received targeted bilateral lesions as shown for sham (open, n=10), mOFC (purple, n=8) or lOFC (green, n=12). (B) Sections showing NeuN staining for sham (left), mOFC (middle) and lOFC (right) lesion groups. (C) Sessions to reach training criteria was not different between groups. (D) The number of trials initiated was not different in the first block, but signficantly reduced in lOFC- compared to mOFC-lesioned rats after acquisition. (E) lOFC-lesioned rats producing significantly fewer correct sequences than sham rats across acquisition. (F) All rats significantly reduced incorrect sequences over acquisition. During early acquisition, lOFC-lesioned rats continued to make more incorrect sequences in block 2 and both lesion groups made more errors in block 3 compared to the sham group. (G) There was a trend for reduced sequence execution time across acquisition, however only the mOFC-lesioned group significantly reduced sequence duration from the first to last block. (H) At the end of acquisition, nose poke duration was faster in lesioned rats than sham controls in the middle of the sequence, however lOFC-lesioned rats were slower at terminating the sequences compared to mOFC-lesioned rats. Data shown as group mean ± S.E.M. *p<0.05.

There was no difference between groups in the first block, but by the end of acquisition the lOFC-lesioned rats initiated fewer trials than mOFC-lesioned rats (Lesion, F_2,27_=4.49, p=0.021; post-hoc comparison p=0.006). lOFC-lesioned rats were also the only group to show a significant *reduction* in trials completed from the first to last block (t _9_=3.17, p=0.011). The number of trials initiated is typically a combination of the reduction in trials as they learn to suppress incorrect sequences and subsequent increase in correct trials as they become more efficient. There was a main effect of Lesion on the number of correct sequences completed (Figure 3E; F_2, 27_=3.55, p=0.043) with lOFC-lesioned rats producing significantly fewer correct sequences than sham treated rats (p=0.014) throughout acquisition. There was also a significant Lesion X Block interaction for the number of incorrect sequences produced (Figure 3F; F_5, 108_=2.59, p=0.034). This was most evident in the early blocks with more errors from lOFC-lesioned rats in block 2 (p=0.026) compared to sham rats. Together, these results show that lOFC-lesioned rats were producing fewer correct and more incorrect sequences, suggesting they were impaired in developing efficiency through invariance and/or learning from negative feedback. mOFC-lesioned rats also made more incorrect sequences in block 3 (compared to sham: mOFC p=0.042, lOFC p=0.055) but did not have other deficits, suggesting this to be a subtle impairment.

#### Sequence timing

There was an overall significant reduction in total sequence duration across acquisition (Figure 3G; F_4, 108_=11.11, p<0.001), however only the mOFC-lesioned group showed a significant reduction in duration from the first to last block (sham: t_7_=1.52, p=0.17; mOFC: t _11_=5.28, p<0.001; lOFC: t_9_=1.14, p=0.29). Rats became significantly faster at executing nose pokes from the first to last block (F_1,27_=26.28, p<0.001) with a significant Block X Hole interaction (Figure 3H; F_4, 108_=22.33, p<0.001) as response times shifted to a ballistic response pattern with training. Between treatment groups, there was a significant Lesion X Hole (F_5,64_ =2.68, p=0.032) interaction with both lesion groups making faster responses in the middle of the sequence than sham rats (hole 3 p’s<0.003), yet lOFC-lesioned rats were significantly delayed on the terminal action in the sequences compared to mOFC- lesioned rats (hole 5 p=0.011). There was no significant change in the duration of time spent in the magazine (Supplementary Figure 2F) or latency to collect the reward (Supplementary Figure 2E) either over training or between groups. The inter-poke intervals were also not significantly different for lesioned rats, although they appeared slower on the first block leading to a significant Block X Lesion interaction following the shift to sham levels by the final block (Supplementary Figure 2A, B). The lOFC-lesioned rats were highly efficient at mid-sequence execution but had relatively elongated terminal nose pokes, when rats usually pause to detect cues associated with pellet delivery and start the next motor plan - reward collection.

In summary, lOFC-lesioned rats were impaired across many measures of action sequence acquisition. While they performed as well as sham rats in the first block, they did not adapt efficiently to the requirement to only produce invariant sequences. This was evidenced by the more gradual reduction in incorrect sequences, consistently fewer correct sequences and start/stop delays observed when initiating and terminating sequences (despite unimpaired mid-sequence execution). Given shared impairments in initiating and automatising sequencing, the lOFC to DLS projection may be important for loading motor sequences. This is in contrasts to mOFC-lesions, which reduced their sequence duration across acquisition, but also produced more incorrect responses during acquisition, unlike the enhancing effects of DMS lesions.

### Prelimbic and infralimbic cortex lesions do not alter sequence acquisition

To further understand the role of the medial prefrontal cortex, we next examined the effects of excitotoxic lesions of the prelimbic (PrL) and infralimbic (IL) cortex. These regions are associated with goal-directed and habitual behavior respectively, with the PrL having strong inputs to the DMS and the IL into the ventral striatum (Coutureau & Killcross, 2003; Hart, Leung, & Balleine, 2014; Heilbronner, Rodriguez-Romaguera, Quirk, Groenewegen, & Haber, 2016; Mailly, Aliane, Groenewegen, Haber, & Deniau, 2013).

#### Acquisition of sequencing

Identical procedures were implemented in PrL and IL lesioned rats (Figure 4A, B). All groups reached criteria before moving onto the sequence acquisition (Figure 4C). The number of correct sequences significantly increased across acquisition (Figure 4E; F_2,51_=26.57, p<0.001) and incorrect sequences significantly decreased (Figure 4F; F_2, 41_=58.93, p<0.001) with no effect of treatment or interactions on trials initiated (Figure 4D) or the number correct or incorrect sequences (Figure 4E, F).

**Figure 4.**
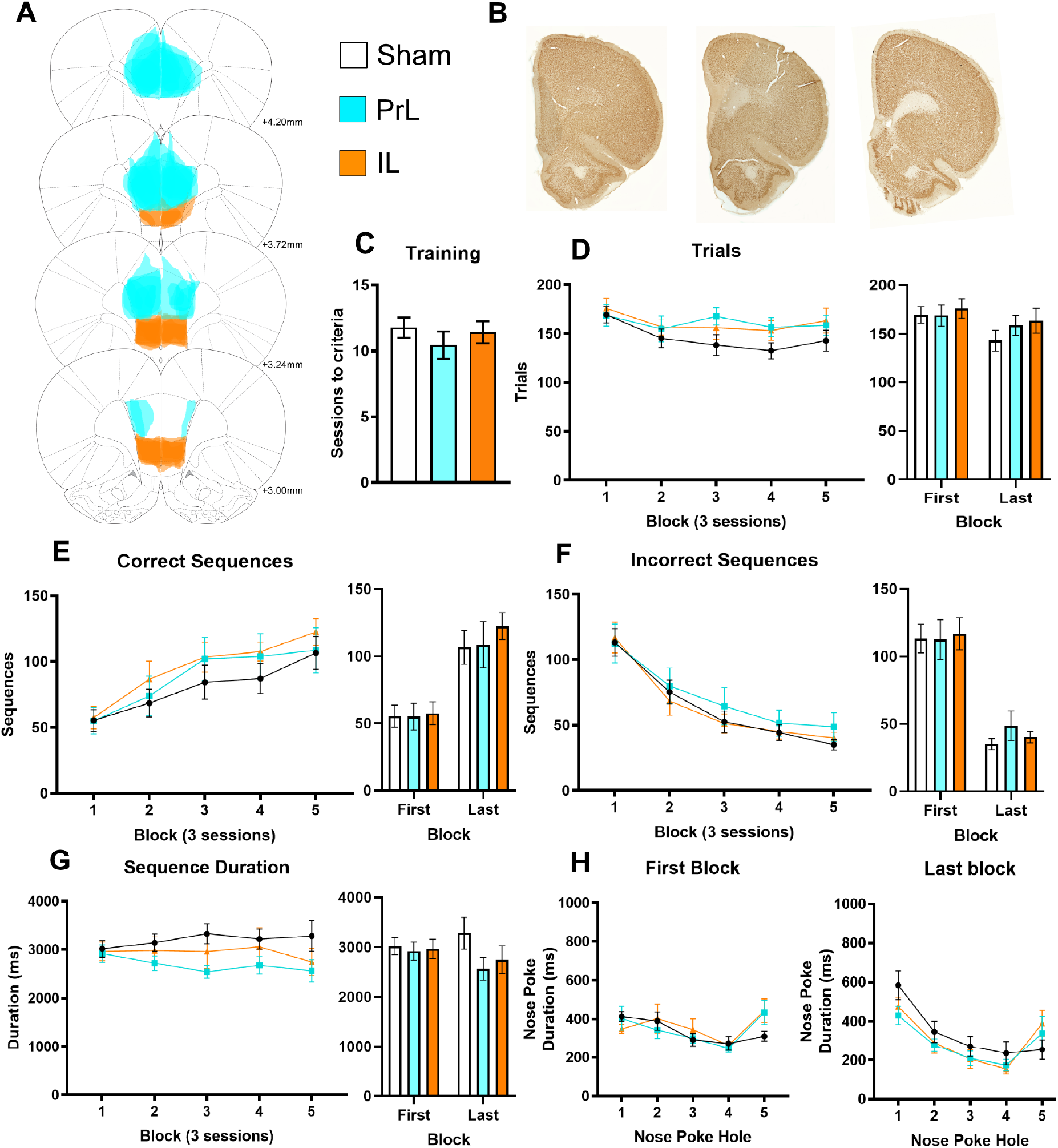
Prelimbic and infralimbic cortex lesions do not alter sequence acquisition. (A) Rats received targeted bilateral lesions as shown for sham (open, n=9), PrL (cyan, n=9) or IL (orange, n=7). (B) Sections showing NeuN staining in sham (left), PrL (middle) and IL (right) lesion groups. (C) The number of trials initiated was not different between groups. (D) The number of correct sequences significantly increased without an effect of lesion. (E) Incorrect sequences significantly decreased, and this was also not different between groups. (F) Rats became significantly faster at executing the sequence with training with no significant differences between groups. (G) Total sequence duration reduced across the acquisition period but was not different between groups. (H) Nose poke duration shifted to the characteristic accelerating pattern with no effect of PrL or IL lesion. Data shown as group mean ± S.E.M. *p<0.05.

#### Sequence timing

While there was a main effect of Block (Figure 4G; F_3, 64_=2.95, p=0.041) on total sequence duration where rats became significantly faster at executing the sequence with training there was no significant difference between lesion groups for nose poke duration across sequence or magazine (Supplementary Fig 3F), inter-poke intervals between holes (Supplementary Fig 3E), or interval from hole 5 to the magazine (Supplementary Fig 3D). Nose poke duration did reduce from first to last block (F_1, 22_=7.61, p=0.011) across all lesion groups and a significant Block X Hole interaction (Figure 4H; F_4, 2_=8.64, p<0.001) identified a ballistic-like response pattern with training. These results indicated that the PrL and IL cortex were not critical for the acquisition of action sequencing.

## DISCUSSION

We found that heterogenous action sequences can come under habitual control, as defined by outcome devaluation insensitivity, when the parameters of this task promoted automaticity. Using this task, we provide the first direct causal evidence that the DMS and DLS have opposing roles on the acquisition of action sequencing. We demonstrated this competitive relationship by showing that DMS lesions enhanced action sequence acquisition and DLS lesions impaired it. The finding of striatal opposition is consistent with studies showing concurrent activity within the DMS and DLS across numerous tasks and training stages (Thorn & Graybiel, 2014). These results also build on recordings in rodents showing that disengagement of the DMS predicts skill learning by allowing the DLS to take control (Kupferschmidt et al., 2017). And that the DMS gates habit formation in the T-maze as although the DLS is active early during learning it only gains control when DMS activity subsides (Thorn et al., 2010). While the lOFC was required for efficient sequencing, surprisingly lesions to medial prefrontal cortical subregions (mOFC, PrL and IL) did not impair nor enhance acquisition of action sequencing. Together, these results demonstrate that reduced DMS activity facilitates the acquisition of DLS-dependent skills and habits, but this is not the product of modulation from cortical inputs. Although we did not investigate the anterior cingulate cortex, our results suggest the source of arbitration between these parallel corticostriatal loops is independent of these prefrontal inputs to the dorsal striatum.

### Opposing roles of the dorsal striatum in the acquisition of action sequences

Previous studies have shown habitual responding can be acquired despite DMS lesions (Gremel & Costa, 2013; Hilario, Holloway, Jin, & Costa, 2012), suggesting a DMS-dependent goal-directed acquisition phase is not required for the development of habits. We provide evidence for this hypothesis by demonstrating not only that DMS-lesioned rats were capable of performing automatised action sequences, but that they show *enhanced* acquisition and reduced variability of this habit-like response pattern. These results not only indicated that DMS-dependent learning was not critical for efficient task acquisition, but that the DMS hampers the development of action sequencing. In contrast, rats with DLS-lesions were impaired in acquiring action sequencing, which is entirely consistent with this task measuring skill formation and sequencing under habitual control.

These results are also consistent with findings from three studies where the inverse pattern (i.e. DMS impairs and DLS enhances performance) was found using tasks that require flexible or goal-directed responding. Moussa, Poucet, Amalric, and Sargolini (2011) found DMS lesions impaired T-maze acquisition, but DLS lesions enhanced learning rate. A second example of striatal opponency was demonstrated by Bradfield and Balleine (2013), where removing the influence of the DLS enhanced goal-directed control beyond the capacity of sham treated rats. A third example comes from a study of visual discrimination, where silencing the DLS during the choice phase led to faster learning, again highlighting that removing DLS activity enhances adaptive behaviors beyond those seen when both regions are functional (Bergstrom et al., 2018). When considered with the results of the current study, these results support a competitive opponency between the DLS and DMS by utilising tasks optimised by flexible responding or automaticity (see Figure 5).

**Figure 5.**
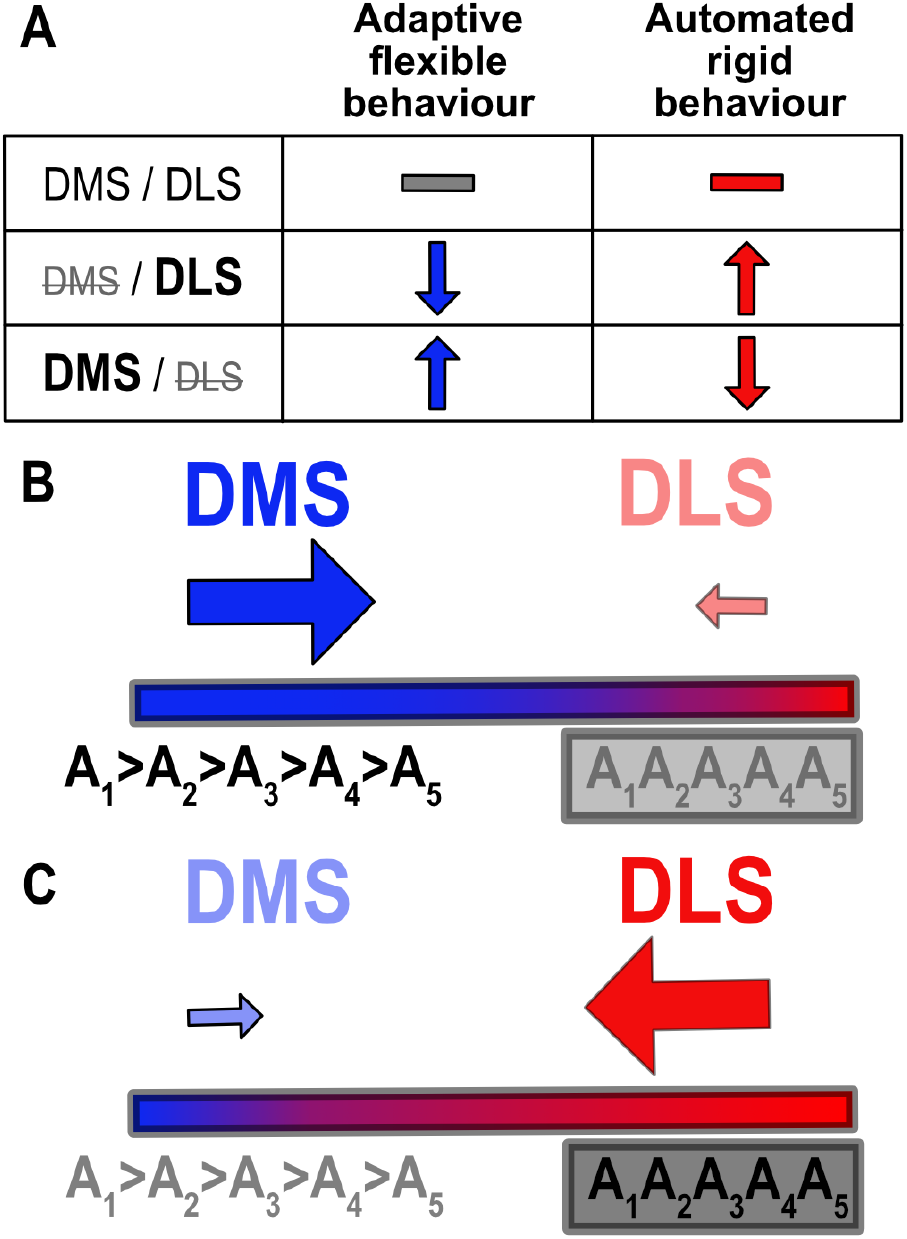
Competitive parallel control by the dorsomedial and dorsolateral striatum. (A) Studies of flexible behavior have found DMS lesions impair performance as anticipated given the role of the DMS in goal-directed behaviors. However, studies have also found DLS-lesioned rodents showed *enhanced* learning compared to controls, suggesting a competitive influence of DLS functions on DMS-dependent behaviors (Bergstrom et al., 2018; Bradfield & Balleine, 2013; Moussa et al., 2011). We build on this model by demonstrating that the converse is true for automatisation of actions. DLS lesions unsurprisingly impaired performance where the task demands habit like behavior. However, we found that DMS lesions enhanced acquisition, suggesting this competitive relationship is bidirectional. (B) Based on these findings we propose a model of opponency between the DMS and DLS. In situations where adaptive or goal-directed behaviors are critical, DMS control dominates and results in performance of individual, slower actions that can be easily modified. Lesioning the DLS biases behavior in this direction. We suggest that just as a purple color gradient can be made bluer through either adding more blue (enhanced DMS activity) or not adding as much red (DLS lesioning); the relative balance is critical such that the loss of one region’s function enhances expression of the other. Parallel development of both pathways incorporates redundancy such that either region can take control as situations change. (C) Tasks requiring automatised actions, such as action sequencing and chunking, occur under DLS-dominated control. Disengagement of the DMS to allow DLS domination has been proposed in the transition from goal-directed to habitual action and in skill refinement (Kupferschmidt et al., 2017). This study demonstrates that habit-like behaviors can also be expedited via DMS loss of function, indicative of functional opponency.

### Habits, skills and automaticity

Here we capitalised on a task that is dependent on reduced behavioral variation (rather than overtraining) to examine the neural underpinnings of automatisation, reflecting the shared features of habits and skills. How the similarities and differences between habits and skills can be consolidated has been a question of growing interest that remains largely unanswered (Ashby et al., 2010; Graybiel & Grafton, 2015; Hardwick et al., 2019; Robbins & Costa, 2017). While acknowledging that each is defined by specific characteristics, these results sit at the intersection of skills and habits and are therefore discussed in this broader context.

Automaticity is commonly measured in skill learning using tasks such as rotarod (Kupferschmidt et al., 2017; Yin et al., 2009) and action sequencing paradigms, including fixed ratio lever pressing or shorter two-step sequencing (e.g. L-R lever press) (Cui et al., 2013; Garr & Delamater, 2019; Jin, Tecuapetla, & Costa, 2014; Tecuapetla, Jin, Lima, & Costa, 2016; Wassum, Ostlund, & Maidment, 2012; Yin, 2009, 2010; Yin, Ostlund, et al., 2005). A four-step (L-L-R-R) lever press task was developed using no experimental cues and a self-paced design (Geddes, Li, & Jin, 2018), however, to our knowledge, models of skill and habit formation have not been tested in rodent operant paradigms requiring more than two different response elements. We found that DLS-lesions specifically affected sequence initiation rather than execution elements, which is in agreement with the suggestion that DLS activity is important when starting and stopping motor sequences, rather than the mid-sequence actions, which is evident in task bracketing patterns within the DLS (Jin & Costa, 2010; K. S. Smith & Graybiel, 2013; Sternberg et al., 1978). It has been suggested that rather than identifying the specific motor actions that will be performed, DLS activity may be important for bracketed groups of familiar motor actions as a chunk (K. S. Smith & Graybiel, 2013). Our results lend support to this suggestion as DLS-lesioned rats did not have deficits in performing the five actions in the correct order (which would be evidenced by an increase in errors) and displayed a ballistic response pattern synonymous with chunking but were impaired when starting the sequence. This has important implications for the role of the DLS in automaticity, habits, and skill formation. Although it is unclear how each concept applies across initiation, execution, and termination elements with action sequences, it is plausible that the DLS is important for retrieving and initiating rehearsed behavioral patterns, promoting their rapid, stimulus-driven and refined expression. Isolating the role of striatal circuits within sequence performance is also critical for understanding movement disorders such as Parkinson’s disease, where action initiation is impaired (Agostino, Berardelli, Formica, Accornero, & Manfredi, 1992).

We demonstrate that heterogenous sequences do lead to habitual responding under certain conditions. Reduced variation through rigid repetition may be a critical condition for the development of habits. We observed this when establishing the task, but also as significantly reduced variation in sequence duration across acquisition in DMS-lesioned rats. Indeed, using an FR5 lever press task Vandaele and Janak (2021) recently reported that rats performed habitually under strict sequencing conditions (DT5), but that allowing rats to either make mid-sequence reward port entries or greater than five presses reverted behavior to goal-directed control. This was accompanied by high DLS and low DMS activity during the DT5 task, but relatively similar activity across the striatum in the task variants. As pointed out by Dickinson (1985), *“contrary to popular belief, habit formation is not a simple consequence of over-training or practice. Rather it appears to arise because over-training typically tends to reduce the variation in behaviour…”* (page 76). Similarly, Daw et al. (2005) suggested that the shift from a model-free to model-based control is dependent on uncertainty, where even providing two choices will prevent model-free responding. Further, Drummond and Niv (2020) suggest that the level of *certainty* within the model-based and model-free estimates may determine which system becomes engaged. A recently proposed dual-system model suggests goal-directed and habitual responding are acquired in parallel, with prediction error determining the associative strength of these processes and responses reflecting their summation (Perez & Dickinson, 2020). This account suggests that as actions become more stereotypical the goal-directed contribution wanes and habitual responding remains. Experimental support for the notion that habits are not merely the product of overtraining was also demonstrated across five human studies that failed to produce habitual responding (de Wit et al., 2018) and by a recent consortium across four laboratories where extended training did not produce habitual responding (Pool et al., 2021). Overtraining also did not elicit habitual responding on a rodent L-R lever pressing task (Garr & Delamater, 2019). In addition, evidence from Hardwick et al. (2019) suggests habits form easily, but their expression can be overruled by goal-directed control such that time to act is also critical factor in determining which is expressed. Action sequencing that is invariant, outcome insensitive and rapid as in this sequential nose poke task provides the ideal platform to examine the neural circuits that support automaticity, habits, and skill formation.

### Cortical functions in action sequencing

Cortical inputs may play a critical role in goal-directed learning, habit formation and skill development but less is known about how they operate across transitions and in action sequences (Bassett, Yang, Wymbs, & Grafton, 2015; Bergstrom et al., 2018; Bradfield et al., 2018; Gremel & Costa, 2013; Killcross & Coutureau, 2003; Kupferschmidt et al., 2017; K. S. Smith & Graybiel, 2013; Turner & Parkes, 2020). A link between cortical disengagement and skill refinement has been observed using imaging in humans (Bassett et al., 2015) and recordings in rodents (Kupferschmidt et al., 2017). As these are correlational findings, reduction in cortical activity may not be critical to skill refinement but may be a consequence of changes in other regions within cortico-striatal loops. Previous research has associated PrL with goal-directed actions and the IL with habits. Using a robust lesioning approach, our results provide the first evidence that these regions are not required to learn and perform heterogenous action sequences.

The PrL cortex is important for early stages of goal-directed learning but not for habit formation (Corbit & Balleine, 2003; Coutureau & Killcross, 2003; Hart, Bradfield, Fok, Chieng, & Balleine, 2018), which is consistent with the lack of effect in this study where goal-directed control was minimised. The fact that PrL lesions did not *enhance* sequencing indicates that the PrL inputs to the DMS are not solely responsible for maintaining DMS functions or goal-directed interference on this task and the role of the PrL cortex is clearly separable. This independence of functions between the PrL cortex and DMS suggests the switch in control within the dorsal striatum is not driven by the PrL cortex.

Lesioning the IL did not impair sequence acquisition as would have been predicted from devaluation studies where IL-lesions result in goal-directed responding (Coutureau & Killcross, 2003). Shipman, Trask, Bouton, and Green (2018) suggested that control shifts from the PrL to IL with experience but prior to habit formation, highlighting a role in the transition of control. Further, K. S. Smith and Graybiel (2013) proposed that the IL and DLS operate together to establish habits, however we found no IL-related deficit in sequence acquisition as was observed for DLS lesions. This suggests that the IL was not required for the automatisation or chunking of action sequences. It is important to note that there are differences between the electrophysiological signatures of DLS and IL in habits (e.g., after devaluation), and there are no direct IL-DLS projections, suggesting they have independent roles in habitual responding. In addition, IL activity does not reflect the habitual nature of individual decisions, indicating it is not arbitrating between goal-directed and habitual strategies but instead reflects overall response tendencies or states (K. S. Smith & Graybiel, 2013). Haddon and Killcross (2011) found that the IL plays a role when goal-directed and habitual associations are in competition, but this was not the case in our study as flexible, goal-directed responding was not advantageous. Our results support the argument that competition, particularly in the context of extended training, may be an important condition for IL-dependent habits (or suppression of goal-directed control), as with little-to-no competition, IL lesions do not influence action sequence acquisition.

In contrast, lOFC lesions reduced total sequences with fewer correct sequences (and increased incorrect sequences) and delayed sequence termination. While largely consistent with deficits in DLS-lesioned rats, two key differences emerged (i) lOFC lesioned rats were relatively slower to terminate sequences and (ii) had higher rates of incorrect responses. The terminal delay in our study, as well as the delayed reward collection latency reported in Hervig et al. (2019), may be due to the lOFC’s role in predicting outcomes based on Pavlovian cues as the reward delivery was cued (Ostlund & Balleine, 2007; Panayi & Killcross, 2014). This is important given the lOFC has been implicated in perseverative and compulsive behaviors, which lack appropriate termination (Burguiere, Monteiro, Feng, & Graybiel, 2013; Chudasama & Robbins, 2003). The lOFC has also been implicated in credit assignment, which is likely to be important when chaining a series of actions where only the final element is followed by reward (Noonan, Chau, Rushworth, & Fellows, 2017). Impaired credit assignment may have diminished learning about more distal sequence elements and increase sequencing errors. The role of the lOFC in using Pavlovian occasion setting cues may also explain the impairment in reducing incorrect responses, which were signalled by the illumination of the house light in this task (Shobe, Bakhurin, Claar, & Masmanidis, 2017).

Prior studies have found that large lOFC lesions produced similar effects to those seen in DMS-lesioned animals performing under both random ratio (RR) and random interval (RI) schedules (Gremel & Costa, 2013). The lack of devaluation sensitivity in both RR and RI contexts following lOFC loss of function was suggested to indicate its role in conveying action-value information. Our results support this notion as an impairment in learning rather than in increase in habit formation, given they made more incorrect responses and performed fewer sequences. Possible roles of the anterior cingulate cortex and motor cortex remain to be tested (Ostlund, Winterbauer, & Balleine, 2009). However, a recent study found that while DLS lesions impaired motor skill performance, motor cortex inputs to the DLS were not required (Dhawale, Wolff, Ko, & Olveczky, 2021). Future studies should confirm if lOFC to DLS projections are critical to action sequencing and isolate the lOFC deficits linked to this specific pathway.

Overall, the cortical effects (or lack thereof) described here are problematic for the popular model of top-down control applied by cortical regions over subcortical structures. This may simply not apply in the same way to behaviors that dominate motor rather than cognitive cortico-striatal loops. This lack of effect is significant in the context of understanding where arbitration of striatal control originates and highlights the importance of considering tasks that optimise automatic, habitual actions to understand cortico-striatal functions. Perhaps when there is little or no need for goal-directed control, there is also little need for medial prefrontal cortical input. However, we also did not observe enhanced acquisition, like the DMS-lesioned rats, which may be due to redundancy within the cortex given multiple sub-regions project to the DMS. We have determined that the medial prefrontal cortex is not responsible for DMS disengagement in skilled, habitual action sequences.

## Conclusions

These findings provide the strongest evidence yet for competition between DMS and DLS functions in the development of behavioral automatisation. We found medial prefrontal subregions were largely unnecessary for sequence acquisition, however lesions to the lOFC impaired action sequencing. Developing an innovative spatial heterogeneous action sequencing task, we were able to isolate initiation, execution and termination specific deficits. These results provide empirical support for a model where DMS activity limits the formation of automated behavior, emphasising its role in gating the acquisition of skills and habits.

## Acknowledgements

NHMRC Early Career Fellowship (GNT1122221) to KMT. This research was funded in whole, or in part, by the Wellcome Trust (Grant 104631/Z/14/Z to TWR). For the purpose of open access, the author has applied a CC BY public copyright licence to any Author Accepted Manuscript version arising from this submission. Experiments were conducted under a Home Office Project Licence held by Dr. Amy Milton and we thank her for supporting our research.

## Author contributions

KMT and TWR designed the experiments; KMT, ML, AS, CM performed the experiments; KMT wrote the first draft and ML, AS, CM, TWR reviewed and edited the manuscript.

## MATERIALS AND METHODS

### EXPERIMENTAL SUBJECT DETAILS

The task was developed in treatment naïve rats where we examined the effects of extended training and then the inclusion of punishment for incorrect sequences. Using this refined protocol, we then conducted three experiments in separate cohorts of rats examining the effect of pre-training lesions of the (1) DMS and DLS; (2) mOFC and lOFC; and (3) PrL and IL on acquisition of action sequencing. Methods were the same across these experiments, with exceptions detailed below.

#### Animals and Housing

Adult male Lister-hooded rats weighing 280-300g (Charles River, UK) were housed in groups of four on reversed 12-h light cycle (off at 07:00) within a temperature (21°C) and humidity-controlled environment in open top cages with aspen bedding, wood block and tube. A week after arriving, rats were food-restricted to no less than 90% of free-feeding weight with unrestricted access to water and were exposed to reward pellets. All procedures were conducted in accordance with the United Kingdom Animal (Scientific Procedures) Act of 1986 and were approved by ethical review at the University of Cambridge.

### METHOD DETAILS

#### Apparatus

Rats were trained to perform a five-step sequential nose poke task (SNT), which was adapted from Keeler, Pretsell, and Robbins (2014), however with substantial changes including absence of cues and the number and order of responses. The task was conducted in operant chambers (Campden Instruments, UK) with five nose poke apertures available within a horizontal array and a reward receptacle on the opposing wall (Robbins, 2002). Nose pokes and the reward receptacle were fitted with infra-red beams to detect head entries and a light for illumination. Reward sucrose pellets (AIN76A, 45mg; TestDiet, UK) were delivered into the receptacle by a pellet dispenser. A house light was mounted on the ceiling and the chamber was contained within a sound attenuating box. Overhead cameras (SpyCameraCCTV, UK) were mounted above each chamber to monitor and record behavior remotely. Whisker Server software and custom programming software was used to operate the chambers and record responses (Cardinal & Aitken, 2010; Keeler et al., 2014).

#### Sequential Nose poke Task (SNT) Protocol

The SNT requires rats to make a nose poke response into each of the five holes from left to right across a horizontal array to receive a food reward. Sessions ran for 30 min unless stated otherwise and all nose pokes and head entries were recorded with the duration of each nose poke calculated based on the entry and exit times. Rats were first habituated to the chambers and retrieved rewards from the receptacle that were dispensed with each head entry until 100 were collected (stage 1). Next, rats were trained to make nose poke responses into the five-hole array (stage 2). Each hole in the five-step sequence was illuminated for 1 s before moving to the next location from left to right and finishing with reward delivery (e.g. 1-2-3-4-5-Reward), which was signalled by illumination of the receptacle. Head entry into the receptacle triggered the start of the next trial. Critically, when the rat nose poked an illuminated hole, the light and sequence counter immediately moved on to the next hole, allowing the rat to achieve reward delivery faster than if they did not nose poke. If the rat made a nose poke into an alternative hole, the illuminated hole would flash for the duration of the incorrect nose poke to draw attention to the correct location. To further encourage nose poking, the illumination duration incremented by 10% of the original delay (1 s) each trial, further delaying reward delivery if nose pokes were not made. This training protocol was implemented to reduce bias for the start or end elements (inherent to training by chaining) and rapidly produced sequencing behavior. Once rats were successfully able to complete at least 15 sequences within a session, they moved to stage 3 where the illumination sequence only advanced to the next hole, and ultimately to reward delivery, after a correct nose poke response into an illuminated hole. Criteria for stage 3 was 50 complete sequences, which was typically achieved in a single session. Stage 4 was identical to stage 3, except that now the nose poke holes were no longer illuminated. After each of the holes had been poked in order, a reward was delivered. Incorrect nose pokes were recorded, but not punished. After reaching 50 uncued sequences, they were moved to the final level (stage 5) where incorrect nose pokes were punished with a 5 s time out period signalled by the illumination of the house light.

After the time out ended, the rat was required to start the sequence again from hole 1. Responses during the timeout period were recorded but did not extend the time out duration. Testing on stage 5 was conducted for 15 sessions and rats began immediately after reaching training criteria. Key measures included trials initiated, correct sequences, incorrect sequences, nose poke durations at each location and total sequence duration.

#### Task development

During task development we originally only trained to stage 4. Rats were then split into two groups (n=12) with one group continuing with daily training sessions (morning only), while the extended group moved to twice daily sessions (morning and afternoon) for 10 days. Sensitivity to outcome-specific devaluation was then tested. As this did not result in habitual action sequencing, rats were then reallocated (matched for prior training history) to either continue daily training sessions at stage 4 (flexible group) or were moved to stage 5 (invariant group) where incorrect sequences were punished for 15 sessions. Rats then underwent outcome-specific devaluation testing.

#### Outcome-specific devaluation

Rats were familiarised to the grain pellets in their home cage prior to devaluation testing. Individuals were placed in empty wire-top cages with free access to 25g of either grain or sucrose pellets for 30 min before being placed into the operant chambers for a 10 min test in extinction. Rats were given two standard training sessions to recover high response rates before being tested with the alternative outcome.

#### Surgery

Prior to training rats were randomly assigned to receive either sham surgery or intracranial bilateral lesions to the region of interest under 2-3% isoflurane anaesthesia with local application of bupivacaine (2mg/kg s.c. at 0.8ml/kg; Sigma) at the incision site. Fibre- sparing lesions were induced by quinolinic acid (0.09M in PBS, Sigma Aldrich, UK) or phosphate-buffered saline (PBS) sham infusions at 0.1ml/min using the co-ordinates in Table 3 relative to bregma based on Paxinos and Watson (2005). Rats were treated with Metacam (1mg/kg; Boehringer Ingelheim) pre- and post-operatively and rehoused in groups of four after lesion surgery. After at least 7 days recovery, rats were food restricted and began operant training as described above.

**Table 1.** Summary of training stages and criteria to move to the next stage.

| Stage | Summary | Criteria | Av. Sessions |
| --- | --- | --- | --- |
| Stage 1 | Habituation to chamber | 100 pellets x 1 session | 1 |
| Stage 2 | Start nose poking 5 holes | >15 sequences x 1 session | 7 |
| Stage 3 | Cued sequence – must NP | >50 sequences x 1 session | 1 |
| Stage 4 | No cues | >50 sequences x 1 session | 3 |
| Stage 5 | Incorrect = Time Out | Final stage | 15 |

**Table 2.** Behavioral measures used to quantify action sequencing.

|  |  |
| --- | --- |
| Trials | Total number of trials initiated |
| Correct | Number of completed sequences |
| Incorrect | Number of incorrect sequences |
| Sequence Duration | NP entry at NP1 to exit on NP5 |
| NP Duration | Time from entry to exit of correct nose poke |
| Inter-Poke Interval (IPI) | Time from exit of previous NP to entry of next NP |
| Initiation Latency | Time from exit magazine to entry NP1 of next trial |
| Reward Latency | Time from exit NP5 to magazine entry when correct |

**Table 3.**
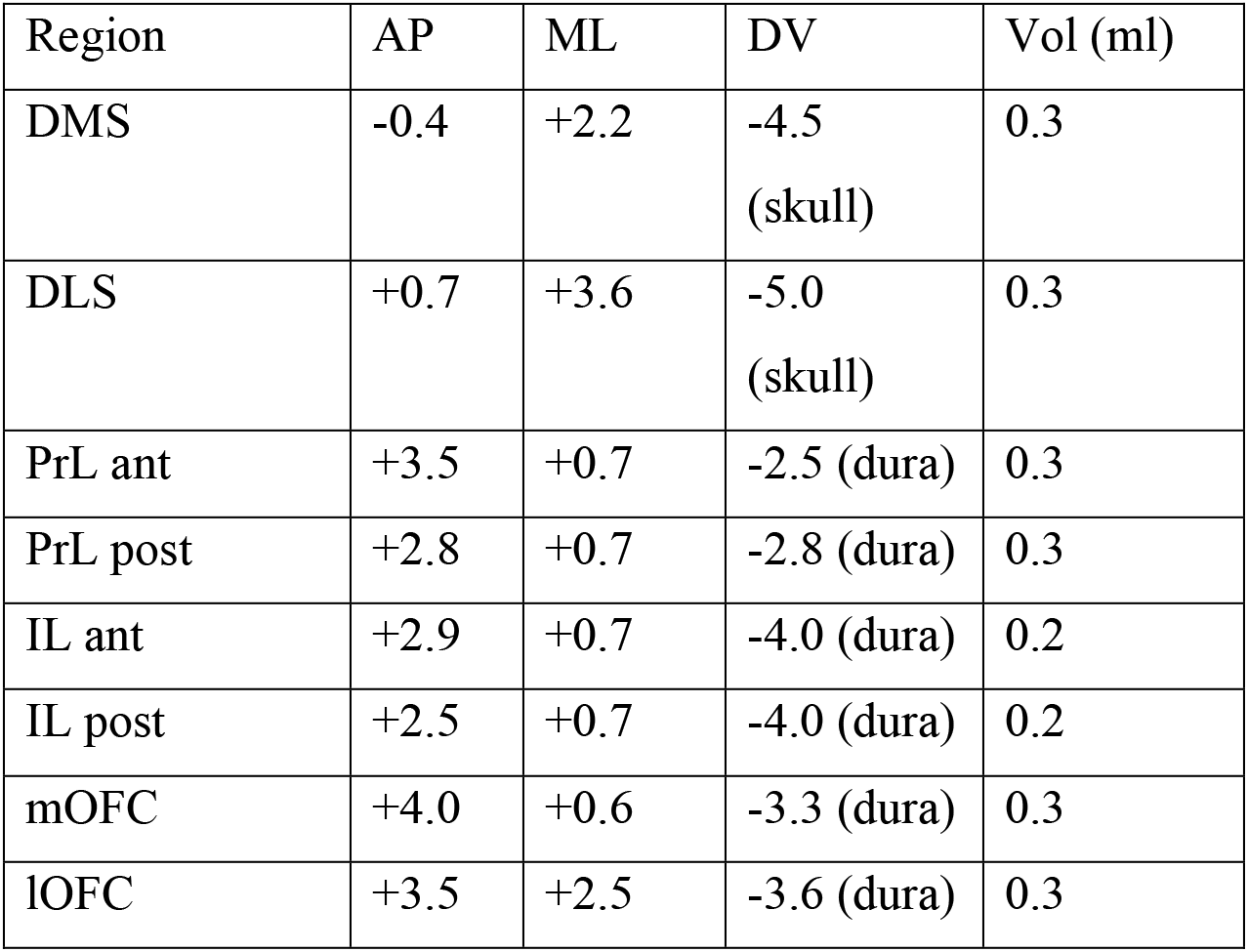
Co-ordinates and volumes used for pre-training lesion infusions of quinolinic acid. DMS: dorsomedial striatum; DLS: dorsolateral striatum; PrL: prelimbic cortex; IL: infralimbic cortex; mOFC: medial orbitofrontal cortex; lOFC: lateral orbitofrontal cortex; ant: anterior; post: posterior.

#### Locomotion

After completion of operant testing, rats were tested for 30 min in an open field arena to rule out gross locomotor impairments. Testing was conducted in lidded boxes (48 x 26.5 x 21cm, Techniplast, UK) in a quiet room with dim red lighting. Locomotion was recorded by infra-red beams across the arena (Photobeam Activity System, San Diego Instruments).

#### Histology

Rats were transcardially perfused using 0.01M PBS with 5g/L sodium nitrite followed by 4% formaldehyde. Brains were then removed for storage in 4% formaldehyde at room temperature overnight on a shaker. They were then transferred to 30% sucrose until they sank before being rapidly frozen and cut into 60mm sections on a freezing microtome (Leica).

Sections were stained for NeuN to confirm lesion placement.

#### NeuN protocol

Sections were washed in 0.01M PBS and then placed in primary antibody (NeuN monoclonal mouse anti-neuronal nuclear protein, Millipore MAB377, 1:2000 in 0.4% Triton X-100 in 0.01M PBS) for two hours on a rotary shaker. Sections are washed three times in 0.01M PBS over 30 min, then secondary (biotinylated anti-mouse IgG, Vector Laboratories BA-2001, at 1:200 in 0.4% Triton X-100 in 0.01M PBS) applied for 90 min. Sections were washed three times in 0.01M PBS, before applying aN immunoperoxidase procedure (Vectastain ABC Kit, Vector Laboratories). Sections were washed three times in 0.01M PBS before visualising in DAB (ImmPACT DAB Peroxidase (HRP) Substrate, Vector Laboratories) and stopping reaction with cold 0.01M PBS. Sections were mounted on gelatin coated slides and dried before clearing with 100% ethanol (2 min), then 50% Ethanol/50% xylene (2 min) and 100% xylene before cover slipping with DPX mountant (Sigma). Images were captured using a NanoZoomer digital slide scanner and visualised with the NDP.view software (Hamamatsu) for histological verification of lesion placement.

### QUANTIFICATION AND STATISTICAL ANALYSIS

#### Statistical Analysis

Rats were excluded for inaccurate or insufficient lesion placement or if they failed to perform action sequences (>20 sessions of training). Final group sizes are reported in the figure legends for each group. Acquisition data was collected over 15 sessions and averaged across blocks of three sessions leading to five blocks. Sequence duration was calculated from the onset of nose poke 1 to the offset of nose poke 5, while the nose poke duration was calculated from entry to exit at each hole. The median and standard deviation for each rat on each day was calculated from individual response times. Timing data was not stored by the program for four rats in one session and therefore their times were averaged across two sessions rather than three for that block to prevent exclusion from the entire dataset. Where appropriate we applied paired t-tests, univariate or repeated measures ANOVA, with simple effects used in the case of significant interactions or post hoc comparisons for effects between treatment groups (SPSS v.25, IBM). Greenhouse-Geisser corrections were made if the sphericity assumption was violated and epsilon was <0.75.

### SUPPLEMENTARY FIGURES

**Supplementary Figure 1.**
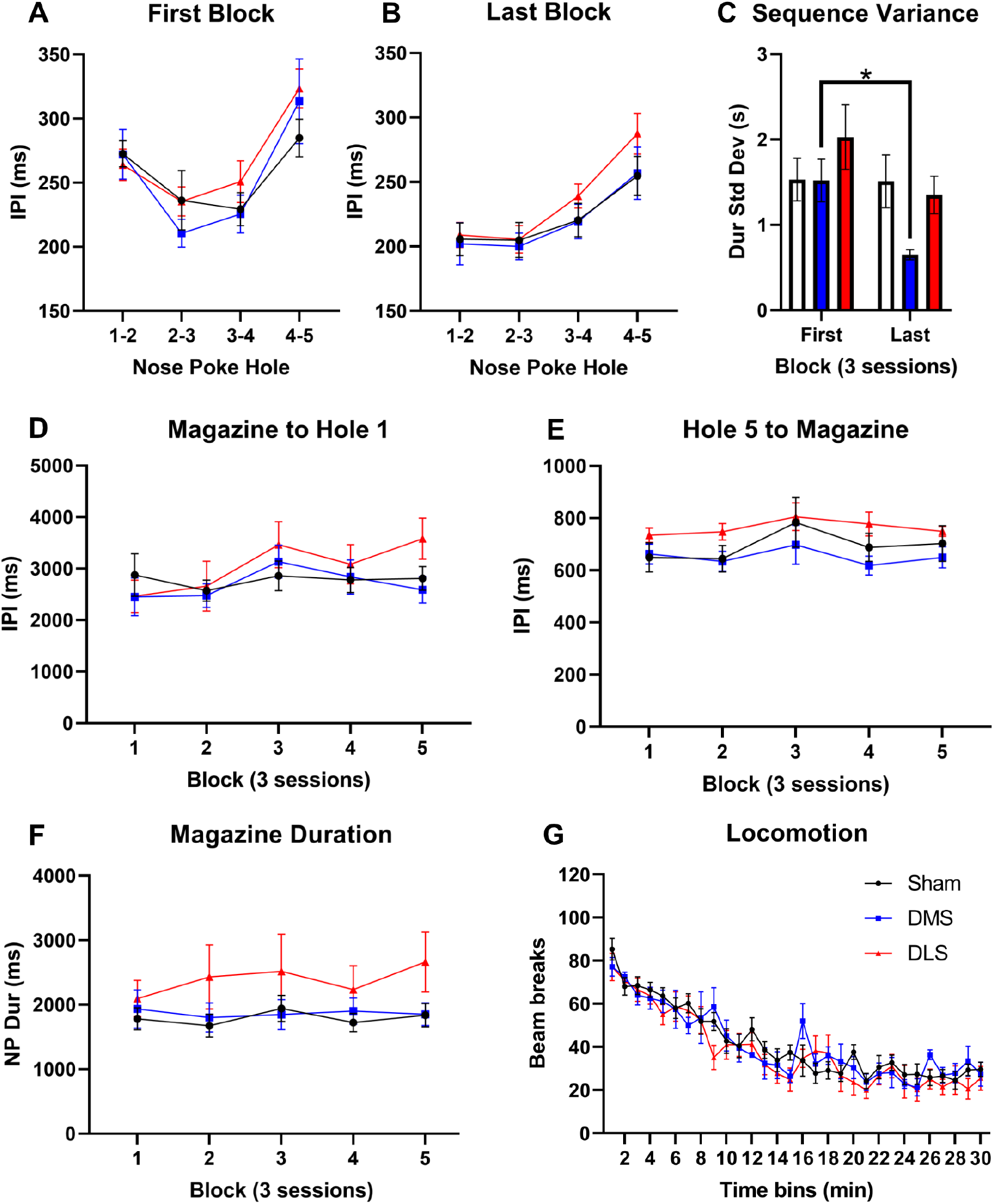
Additional sequencing measures in DMS and DLS lesioned rats. (A) Groups did not differ in the inter-poke interval (IPI) between holes on the first block; sham (open, n=11), DMS lesioned (blue, n=7), DLS lesioned (red, n=8). (B) This remained the case on the last block of acquisition with IPI’s becoming faster with training (Block: F_4, 48_=15.62, p<0.001). (C) There was a significant reduction in the standard deviation of sequence durations, indicating reduced variation with training in the DMS-lesioned rats but not in sham or DLS-lesioned rats (sham: t_10_=0.06, p=0.953; DMS: t_6_=3.09, p=0.021; DLS: t_7_=1.57, p=0.160). (D) The interval between leaving the magazine and nose poking into hole 1 did not differ between groups across acquisition (F_2,23_=0.49, p=0.62). (E) Nor did the interval from the fifth hole of the sequence and magazine entry (reward collection latency; Block F_4,92_=1.91, p=0.16; Lesion F_2,23_=1.46, p=0.25). (F) The time spent with their nose in the magazine also did not significantly differ between groups (Block, F_4,92_=1.23, p=0.30; Lesion, F_2,23_=1.47, p=0.25). (G) There was a main effect of time on locomotor activity, but no effect of treatment (p>0.4).

**Supplementary Figure 2.**
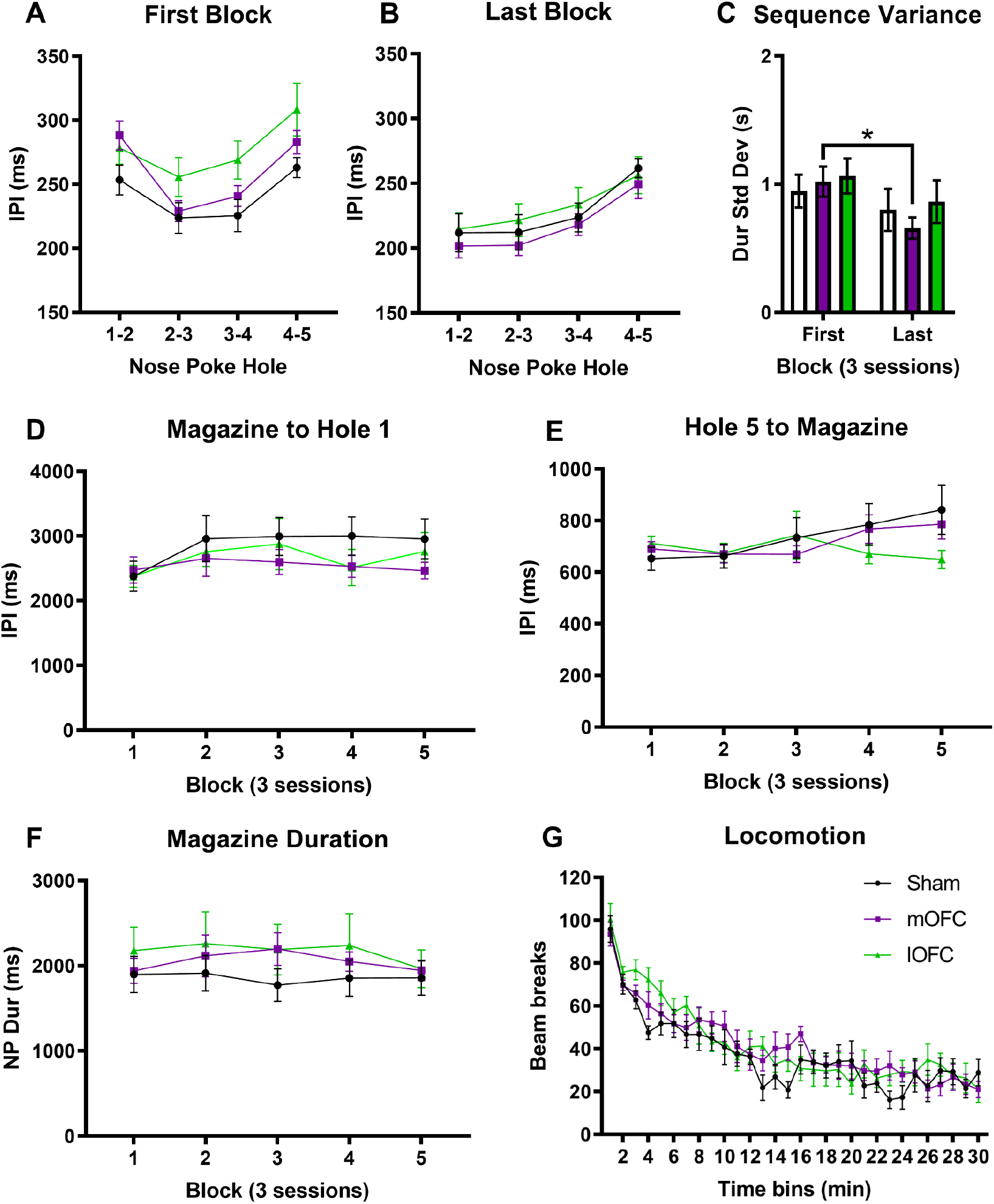
Additional sequencing measures in mOFC and lOFC lesioned rats. (A) Groups did not differ in the inter-poke interval (IPI) between holes on the first block; sham (open, n=10), mOFC (purple, n=8) or lOFC (green, n=12). (B) This remained the case on the last block of acquisition with IPI’s becoming faster with training. However, a significant Block X Lesion interaction highlighted that the lesion groups showed greater reduction in IPI times across acquisition due to the relatively slower IPI times in the first block (Block: F_1, 27_=55.0, p<0.001; Lesion: F_12 27_=1.21, p=0.31; Block X Lesion: F_2, 27_=4.34, p=0.023; Hole and Hole X Block: p<0.001; Hole X Lesion p>0.5; Hole X Block X Lesion: F_6, 51_=2.36, p=0.068). (C) There was a significant reduction in the standard deviation of sequence durations, indicating reduced variation with training in the mOFC-lesioned rats but not in sham or lOFC-lesioned rats (sham: t_7_=0.67, p=0.53; mOFC: t_11_=2.53, p=0.028; lOFC: t_9_=1.31, p=0.22). (D) The interval between leaving the magazine and nose poking into hole 1 did not differ between groups across acquisition. (E) Nor did the interval from the fifth hole of the sequence and magazine entry (reward collection latency; Block: F_2,57_=2.95, p=0.06; Lesion: F_2,27_=0.26, p=0.77; Block X Lesion: F_4,57_=2.35, p=0.06). (F) The time spent with their nose in the magazine also did not significantly differ between groups (Block: F_3,75_=1.07, p=0.37; Lesion: F_2,27_=0.46, p=0.64; Block X Lesion: F_6,75_=0.66, p=0.67). (G) There was a main effect of time on locomotor activity, but no effect of treatment (p>0.4).

**Supplementary Figure 3.**
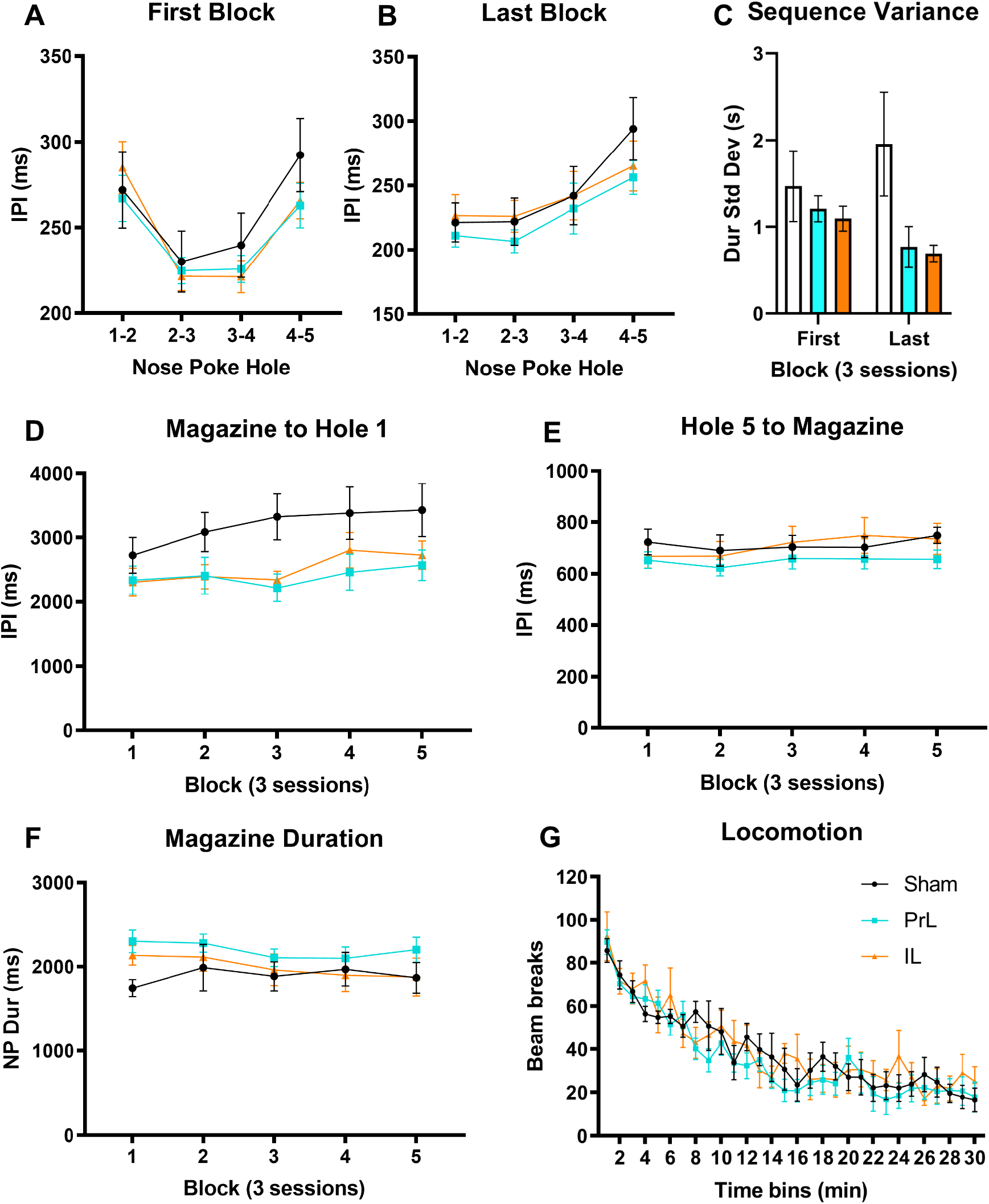
Additional sequencing measures in PrL and IL lesioned rats. (A) Groups did not differ in the inter-poke interval (IPI) between holes on the first block; sham (open, n=9), PrL (cyan, n=9) or IL (orange, n=7). (B) This remained the case on the last block of acquisition. (C) There was a trend towards a reduction in the standard deviation of sequence durations in the IL-lesioned rats but not in sham or PrL-lesioned rats (sham: t_8_=-0.66, p=0.53; PrL: t_8_=1.61, p=0.15; IL: t_6_=2.45, p=0.050). It was noted that two sham rats had rare but excessively long sequence durations, perhaps due to stopping and starting sequencing. Typically, rats would subsequently make an incorrect response if they paused, however here they were still able tom complete a correct sequence and that data is captured in the large error bars for sham rat within both blocks. (D) The effect of lesion on the interval between the magazine and hole 1 neared significance with the sham rats taking longer than the lesioned groups (Block: F_2,52_=3.17, p=0.043; Lesion: F_2,22_=3.34, p=0.054; Block X Lesion: F_5,52_=0.70, p=0.61). (E) Nor did the interval from the fifth hole of the sequence and magazine entry (reward collection latency; Block: F_2,51_=2.69, p=0.07; Lesion: F_2,22_=0.73, p=0.49; Block X Lesion: F_5,51_=0.96, p=0.45). (F) The time spent with their nose in the magazine also did not significantly differ between groups. (G) There was a main effect of time on locomotor activity, but no effect of treatment (p>0.4).

## REFERENCES

Abrahamse, E. L., Ruitenberg, M. F., de Kleine, E., & Verwey, W. B. (2013). Control of automated behavior: insights from the discrete sequence production task. Front Hum Neurosci, 7, 82. doi:10.3389/fnhum.2013.00082

Agostino, R., Berardelli, A., Formica, A., Accornero, N., & Manfredi, M. (1992). Sequential arm movements in patients with Parkinson’s disease, Huntington’s disease and dystonia. Brain, 115 ( *Pt 5*), 1481–1495. doi:10.1093/brain/115.5.1481

Ashby, F. G., Turner, B. O., & Horvitz, J. C. (2010). Cortical and basal ganglia contributions to habit learning and automaticity. Trends Cogn Sci, 14(5), 208–215. doi:10.1016/j.tics.2010.02.001

Balleine, B. W. (2019). The Meaning of Behavior: Discriminating Reflex and Volition in the Brain. Neuron, 104(1), 47–62. doi:10.1016/j.neuron.2019.09.024

Balleine, B. W., & Dezfouli, A. (2019). Hierarchical Action Control: Adaptive Collaboration Between Actions and Habits. Front Psychol, 10, 2735. doi:10.3389/fpsyg.2019.02735

Balleine, B. W., Liljeholm, M., & Ostlund, S. B. (2009). The integrative function of the basal ganglia in instrumental conditioning. Behav Brain Res, 199(1), 43–52. doi:10.1016/j.bbr.2008.10.034

Bassett, D. S., Yang, M., Wymbs, N. F., & Grafton, S. T. (2015). Learning-induced autonomy of sensorimotor systems. Nat Neurosci, 18(5), 744–751. doi:10.1038/nn.3993

Bergstrom, H. C., Lipkin, A. M., Lieberman, A. G., Pinard, C. R., Gunduz-Cinar, O., Brockway, E. T., … Holmes, A. (2018). Dorsolateral Striatum Engagement Interferes with Early Discrimination Learning. Cell Rep, 23(8), 2264–2272. doi:10.1016/j.celrep.2018.04.081

Bradfield, L. A., & Balleine, B. W. (2013). Hierarchical and binary associations compete for behavioral control during instrumental biconditional discrimination. J Exp Psychol Anim Behav Process, 39(1), 2–13. doi:10.1037/a0030941

Bradfield, L. A., Dezfouli, A., van Holstein, M., Chieng, B., & Balleine, B. W. (2015). Medial Orbitofrontal Cortex Mediates Outcome Retrieval in Partially Observable Task Situations. Neuron, 88(6), 1268–1280. doi:10.1016/j.neuron.2015.10.044

Bradfield, L. A., Hart, G., & Balleine, B. W. (2018). Inferring action-dependent outcome representations depends on anterior but not posterior medial orbitofrontal cortex. Neurobiol Learn Mem, 155, 463–473. doi:10.1016/j.nlm.2018.09.008

Burguiere, E., Monteiro, P., Feng, G., & Graybiel, A. M. (2013). Optogenetic stimulation of lateral orbitofronto-striatal pathway suppresses compulsive behaviors. Science, 340(6137), 1243–1246. doi:10.1126/science.1232380

Cardinal, R. N., & Aitken, M. R. (2010). Whisker: a client-server high-performance multimedia research control system. Behavior Research Methods, 42(4), 1059–1071. doi:10.3758/BRM.42.4.1059

Carli, M., Robbins, T. W., Evenden, J. L., & Everitt, B. J. (1983). Effects of lesions to ascending noradrenergic neurones on performance of a 5-choice serial reaction task in rats; implications for theories of dorsal noradrenergic bundle function based on selective attention and arousal. Behav Brain Res, 9(3), 361–380. Retrieved from https://www.ncbi.nlm.nih.gov/pubmed/6639741

Chudasama, Y., & Robbins, T. W. (2003). Dissociable contributions of the orbitofrontal and infralimbic cortex to pavlovian autoshaping and discrimination reversal learning: further evidence for the functional heterogeneity of the rodent frontal cortex. J Neurosci, 23(25), 8771–8780. Retrieved from https://www.ncbi.nlm.nih.gov/pubmed/14507977

Corbit, L. H., & Balleine, B. W. (2003). The role of prelimbic cortex in instrumental conditioning. Behav Brain Res, 146(1-2), 145–157. doi:10.1016/j.bbr.2003.09.023

Coutureau, E., & Killcross, S. (2003). Inactivation of the infralimbic prefrontal cortex reinstates goal-directed responding in overtrained rats. Behav Brain Res, 146(1-2), 167–174. doi:10.1016/j.bbr.2003.09.025

Cui, G., Jun, S. B., Jin, X., Pham, M. D., Vogel, S. S., Lovinger, D. M., & Costa, R. M. (2013). Concurrent activation of striatal direct and indirect pathways during action initiation. Nature, 494(7436), 238–242. doi:10.1038/nature11846

Daw, N. D., Niv, Y., & Dayan, P. (2005). Uncertainty-based competition between prefrontal and dorsolateral striatal systems for behavioral control. Nat Neurosci, 8(12), 1704–1711. doi:10.1038/nn1560

de Wit, S., Kindt, M., Knot, S. L., Verhoeven, A. A. C., Robbins, T. W., Gasull-Camos, J., … Gillan, C. M. (2018). Shifting the balance between goals and habits: Five failures in experimental habit induction. J Exp Psychol Gen, 147(7), 1043–1065. doi:10.1037/xge0000402

Dezfouli, A., & Balleine, B. W. (2012). Habits, action sequences and reinforcement learning. Eur J Neurosci, 35(7), 1036–1051. doi:10.1111/j.1460-9568.2012.08050.x

Dezfouli, A., Lingawi, N. W., & Balleine, B. W. (2014). Habits as action sequences: hierarchical action control and changes in outcome value. Philos Trans R Soc Lond B Biol Sci, 369(1655). doi:10.1098/rstb.2013.0482

Dhawale, A. K., Wolff, S. B. E., Ko, R., & Olveczky, B. P. (2021). The basal ganglia control the detailed kinematics of learned motor skills. Nat Neurosci. doi:10.1038/s41593-021-00889-3

Dickinson, A. (1985). Actions and habits-the development of behavioural autonomy. Philos Trans R Soc Lond B Biol Sci, 308, 67–78.

Drummond, N., & Niv, Y. (2020). Model-based decision making and model-free learning. Curr Biol, 30(15), R860–R865. doi:10.1016/j.cub.2020.06.051

Garr, E., & Delamater, A. R. (2019). Exploring the relationship between actions, habits, and automaticity in an action sequence task. Learn Mem, 26(4), 128–132. doi:10.1101/lm.048645.118

Geddes, C. E., Li, H., & Jin, X. (2018). Optogenetic Editing Reveals the Hierarchical Organization of Learned Action Sequences. Cell, 174(1), 32–43 e15. doi:10.1016/j.cell.2018.06.012

Graybiel, A. M., & Grafton, S. T. (2015). The striatum: where skills and habits meet. Cold Spring Harbor Perspectives in Biology, 7(8), a021691. doi:10.1101/cshperspect.a021691

Gremel, C. M., Chancey, J. H., Atwood, B. K., Luo, G., Neve, R., Ramakrishnan, C., … Costa, R. M. (2016). Endocannabinoid Modulation of Orbitostriatal Circuits Gates Habit Formation. Neuron, 90(6), 1312–1324. doi:10.1016/j.neuron.2016.04.043

Gremel, C. M., & Costa, R. M. (2013). Orbitofrontal and striatal circuits dynamically encode the shift between goal-directed and habitual actions. Nature Communications, 4, 2264. doi:10.1038/ncomms3264

Haddon, J. E., & Killcross, S. (2011). Inactivation of the infralimbic prefrontal cortex in rats reduces the influence of inappropriate habitual responding in a response-conflict task. Neuroscience, 199(0), 205–212. doi:10.1016/j.neuroscience.2011.09.065

Hardwick, R. M., Forrence, A. D., Krakauer, J. W., & Haith, A. M. (2019). Time-dependent competition between goal-directed and habitual response preparation. Nat Hum Behav, 3(12), 1252–1262. doi:10.1038/s41562-019-0725-0

Hart, G., Bradfield, L. A., Fok, S. Y., Chieng, B., & Balleine, B. W. (2018). The Bilateral Prefronto-striatal Pathway Is Necessary for Learning New Goal-Directed Actions. Curr Biol, 28(14), 2218–2229 e2217. doi:10.1016/j.cub.2018.05.028

Hart, G., Leung, B. K., & Balleine, B. W. (2014). Dorsal and ventral streams: the distinct role of striatal subregions in the acquisition and performance of goal-directed actions. Neurobiol Learn Mem, 108, 104–118. doi:10.1016/j.nlm.2013.11.003

Heilbronner, S. R., Rodriguez-Romaguera, J., Quirk, G. J., Groenewegen, H. J., & Haber, S. N. (2016). Circuit-Based Corticostriatal Homologies Between Rat and Primate. Biol Psychiatry, 80(7), 509–521. doi:10.1016/j.biopsych.2016.05.012

Hervig, M. E., Fiddian, L., Piilgaard, L., Bozic, T., Blanco-Pozo, M., Knudsen, C., … Robbins, T. W. (2019). Dissociable and Paradoxical Roles of Rat Medial and Lateral Orbitofrontal Cortex in Visual Serial Reversal Learning. Cereb Cortex. doi:10.1093/cercor/bhz144

Hilario, M., Holloway, T., Jin, X., & Costa, R. M. (2012). Different dorsal striatum circuits mediate action discrimination and action generalization. Eur J Neurosci, 35(7), 1105–1114. doi:10.1111/j.1460-9568.2012.08073.x

Izquierdo, A. (2017). Functional Heterogeneity within Rat Orbitofrontal Cortex in Reward Learning and Decision Making. J Neurosci, 37(44), 10529–10540. doi:10.1523/JNEUROSCI.1678-17.2017

Jin, X., & Costa, R. M. (2010). Start/stop signals emerge in nigrostriatal circuits during sequence learning. Nature, 466(7305), 457–462. doi:10.1038/nature09263

Jin, X., & Costa, R. M. (2015). Shaping action sequences in basal ganglia circuits. Curr Opin Neurobiol, 33, 188–196. doi:10.1016/j.conb.2015.06.011

Jin, X., Tecuapetla, F., & Costa, R. M. (2014). Basal ganglia subcircuits distinctively encode the parsing and concatenation of action sequences. Nat Neurosci, 17(3), 423–430. doi:10.1038/nn.3632

Keeler, J. F., Pretsell, D. O., & Robbins, T. W. (2014). Functional implications of dopamine D1 vs. D2 receptors: A ’prepare and select’ model of the striatal direct vs. indirect pathways. Neuroscience, 282, 156–175. doi:10.1016/j.neuroscience.2014.07.021

Killcross, S., & Coutureau, E. (2003). Coordination of actions and habits in the medial prefrontal cortex of rats. Cereb Cortex, 13(4), 400–408. doi:10.1093/cercor/13.4.400

Kupferschmidt, D. A., Juczewski, K., Cui, G., Johnson, K. A., & Lovinger, D. M. (2017). Parallel, but Dissociable, Processing in Discrete Corticostriatal Inputs Encodes Skill Learning. Neuron, 96(2), 476–489 e475. doi:10.1016/j.neuron.2017.09.040

Lehericy, S., Benali, H., Van de Moortele, P. F., Pelegrini-Issac, M., Waechter, T., Ugurbil, K., & Doyon, J. (2005). Distinct basal ganglia territories are engaged in early and advanced motor sequence learning. Proc Natl Acad Sci U S A, 102(35), 12566–12571. doi:10.1073/pnas.0502762102

Mailly, P., Aliane, V., Groenewegen, H. J., Haber, S. N., & Deniau, J. M. (2013). The rat prefrontostriatal system analyzed in 3D: evidence for multiple interacting functional units. J Neurosci, 33(13), 5718–5727. doi:10.1523/JNEUROSCI.5248-12.2013

Miyachi, S., Hikosaka, O., & Lu, X. (2002). Differential activation of monkey striatal neurons in the early and late stages of procedural learning. Exp Brain Res, 146(1), 122–126. doi:10.1007/s00221-002-1213-7

Moussa, R., Poucet, B., Amalric, M., & Sargolini, F. (2011). Contributions of dorsal striatal subregions to spatial alternation behavior. Learn Mem, 18(7), 444–451. doi:10.1101/lm.2123811

Noonan, M. P., Chau, B. K. H., Rushworth, M. F. S., & Fellows, L. K. (2017). Contrasting Effects of Medial and Lateral Orbitofrontal Cortex Lesions on Credit Assignment and Decision-Making in Humans. J Neurosci, 37(29), 7023–7035. doi:10.1523/JNEUROSCI.0692-17.2017

Ostlund, S. B., & Balleine, B. W. (2007). Orbitofrontal cortex mediates outcome encoding in Pavlovian but not instrumental conditioning. J Neurosci, 27(18), 4819–4825. doi:10.1523/JNEUROSCI.5443-06.2007

Ostlund, S. B., Winterbauer, N. E., & Balleine, B. W. (2009). Evidence of action sequence chunking in goal-directed instrumental conditioning and its dependence on the dorsomedial prefrontal cortex. J Neurosci, 29(25), 8280–8287. doi:10.1523/JNEUROSCI.1176-09.2009

Panayi, M. C., & Killcross, S. (2014). Orbitofrontal cortex inactivation impairs between- but not within-session Pavlovian extinction: an associative analysis. Neurobiol Learn Mem, 108, 78–87. doi:10.1016/j.nlm.2013.08.002

Paxinos, G., & Watson, C. (2005). The Rat Brain in Stereotaxic Coordinates (5th ed.). San Diego: Academic Press.

Peak, J., Hart, G., & Balleine, B. W. (2019). From learning to action: the integration of dorsal striatal input and output pathways in instrumental conditioning. Eur J Neurosci, 49(5), 658–671. doi:10.1111/ejn.13964

Perez, O. D., & Dickinson, A. (2020). A theory of actions and habits: The interaction of rate correlation and contiguity systems in free-operant behavior. Psychological Review, 127(6), 945–971. doi:10.1037/rev0000201

Pool, E., Gera, R., Fransen, A., Perez, O. D., Cremer, A., Aleksic, M., … O’Doherty, J. P. (2021). Determining the Effects of Training Duration on the Behavioral Expression of Habitual Control in Humans: A Multi-laboratory Investigati. doi:10.31234/osf.io/z756h

Robbins, T. W. (2002). The 5-choice serial reaction time task: behavioural pharmacology and functional neurochemistry. Psychopharmacology (Berl), 163(3-4), 362–380. doi:10.1007/s00213-002-1154-7

Robbins, T. W., & Costa, R. M. (2017). Habits. Curr Biol, 27(22), R1200–R1206. doi:10.1016/j.cub.2017.09.060

Schreiner, D. C., Renteria, R., & Gremel, C. M. (2020). Fractionating the all-or-nothing definition of goal-directed and habitual decision-making. J Neurosci Res, 98(6), 998–1006. doi:10.1002/jnr.24545

Shipman, M. L., Trask, S., Bouton, M. E., & Green, J. T. (2018). Inactivation of prelimbic and infralimbic cortex respectively affects minimally-trained and extensively-trained goal-directed actions. Neurobiol Learn Mem, 155, 164–172. doi:10.1016/j.nlm.2018.07.010

Shobe, J. L., Bakhurin, K. I., Claar, L. D., & Masmanidis, S. C. (2017). Selective Modulation of Orbitofrontal Network Activity during Negative Occasion Setting. J Neurosci, 37(39), 9415–9423. doi:10.1523/JNEUROSCI.0572-17.2017

Smith, A. C. W., Jonkman, S., Difeliceantonio, A. G., O’Connor, R. M., Ghoshal, S., Romano, M. F., … Kenny, P. J. (2021). Opposing roles for striatonigral and striatopallidal neurons in dorsolateral striatum in consolidating new instrumental actions. Nature Communications, 12(1), 5121. doi:10.1038/s41467-021-25460-3

Smith, K. S., & Graybiel, A. M. (2013). A dual operator view of habitual behavior reflecting cortical and striatal dynamics. Neuron, 79(2), 361–374. doi:10.1016/j.neuron.2013.05.038

Smith, K. S., & Graybiel, A. M. (2016). Habit formation coincides with shifts in reinforcement representations in the sensorimotor striatum. J Neurophysiol, 115(3), 1487–1498. doi:10.1152/jn.00925.2015

Sternberg, S., Monsell, S., Knoll, R. L., & Wright, C. E. (1978). The Latency and Duration of Rapid Movement Sequences: Comparisons of Speech and Typewriting. In G. E. Stelmach (Ed.), Information Processing in Motor Control and Learning (pp. 117–152): Academic Press.

Tecuapetla, F., Jin, X., Lima, S. Q., & Costa, R. M. (2016). Complementary Contributions of Striatal Projection Pathways to Action Initiation and Execution. Cell, 166(3), 703–715. doi:10.1016/j.cell.2016.06.032

Thorn, C. A., Atallah, H., Howe, M., & Graybiel, A. M. (2010). Differential dynamics of activity changes in dorsolateral and dorsomedial striatal loops during learning. Neuron, 66(5), 781–795. doi:10.1016/j.neuron.2010.04.036

Thorn, C. A., & Graybiel, A. M. (2014). Differential entrainment and learning-related dynamics of spike and local field potential activity in the sensorimotor and associative striatum. J Neurosci, 34(8), 2845–2859. doi:10.1523/JNEUROSCI.1782-13.2014

Turner, K. M., & Parkes, S. L. (2020). Prefrontal regulation of behavioural control: Evidence from learning theory and translational approaches in rodents. Neurosci Biobehav Rev, 118, 27–41. doi:10.1016/j.neubiorev.2020.07.010

Vandaele, Y., & Janak, P. H. (2021). Unveiling the neural correlates of habit in the dorsal striatum. bioRxiv, 2021.2004.2003.438314. doi:10.1101/2021.04.03.438314

Vandaele, Y., Mahajan, N. R., Ottenheimer, D. J., Richard, J. M., Mysore, S. P., & Janak, P. H. (2019). Distinct recruitment of dorsomedial and dorsolateral striatum erodes with extended training. Elife, 8. doi:10.7554/eLife.49536

Wassum, K. M., Ostlund, S. B., & Maidment, N. T. (2012). Phasic mesolimbic dopamine signaling precedes and predicts performance of a self-initiated action sequence task. Biol Psychiatry, 71(10), 846–854. doi:10.1016/j.biopsych.2011.12.019

Yin, H. H. (2009). The role of the murine motor cortex in action duration and order. Frontiers in Integrative Neuroscience, 3, 23. doi:10.3389/neuro.07.023.2009

Yin, H. H. (2010). The sensorimotor striatum is necessary for serial order learning. J Neurosci, 30(44), 14719–14723. doi:10.1523/JNEUROSCI.3989-10.2010

Yin, H. H., & Knowlton, B. J. (2006). The role of the basal ganglia in habit formation. Nat Rev Neurosci, 7(6), 464–476. doi:10.1038/nrn1919

Yin, H. H., Knowlton, B. J., & Balleine, B. W. (2004). Lesions of dorsolateral striatum preserve outcome expectancy but disrupt habit formation in instrumental learning. Eur J Neurosci, 19(1), 181–189. doi:10.1111/j.1460-9568.2004.03095.x

Yin, H. H., Knowlton, B. J., & Balleine, B. W. (2005). Blockade of NMDA receptors in the dorsomedial striatum prevents action-outcome learning in instrumental conditioning. Eur J Neurosci, 22(2), 505–512. doi:10.1111/j.1460-9568.2005.04219.x

Yin, H. H., Knowlton, B. J., & Balleine, B. W. (2006). Inactivation of dorsolateral striatum enhances sensitivity to changes in the action-outcome contingency in instrumental conditioning. Behav Brain Res, 166(2), 189–196. doi:10.1016/j.bbr.2005.07.012

Yin, H. H., Mulcare, S. P., Hilario, M. R., Clouse, E., Holloway, T., Davis, M. I., … Costa, R. M. (2009). Dynamic reorganization of striatal circuits during the acquisition and consolidation of a skill. Nat Neurosci, 12(3), 333–341. doi:10.1038/nn.2261

Yin, H. H., Ostlund, S. B., Knowlton, B. J., & Balleine, B. W. (2005). The role of the dorsomedial striatum in instrumental conditioning. Eur J Neurosci, 22(2), 513–523. doi:10.1111/j.1460-9568.2005.04218.x

